# Selectively increased autofluorescence at certain locations of skin may become a novel diagnostic biomarker for lung cancer

**DOI:** 10.1101/315440

**Authors:** Mingchao Zhang, Yue Tao, Qing Chang, Yujia Li, Tianqing Chu, Weihai Ying

## Abstract

Early diagnosis is critical for improving the 5-year survival rate of lung cancer patients. Our current study tested our hypothesis that increased autofluorescence (AF) of skin and nails may become a novel diagnostic biomarker of lung cancer, which has generated the following findings: First, our study on a mouse model of lung cancer has shown that development of lung cancer led to a marked increase in the epidermal green AF of the mice. Second, the AF intensity of the untreated lung cancer patients was significantly higher than that of the healthy persons and the pulmonary infection patients at certain examined locations of the skin and fingernails. Third, the ‘Pattern of AF’ of healthy controls, pulmonary infection patients and untreated lung cancer patients was markedly different from each other. Fourth, when the number of the locations with increased AF was used as the sole diagnostic parameter, our ROC analysis showed that the AUC was 0.9067 for differentiating the healthy controls and the untreated lung cancer patients. Collectively, our study has indicated that development of lung cancer is sufficient to induce increases in the epidermal green AF of both mice and human subjects. Our study has also indicated that the ‘Pattern of AF’ of lung cancer patients could become a novel biomarker of lung cancer, which holds great promise for non-invasive, rapid and economic diagnosis and screening of lung cancer.

## Introduction

Lung cancer is the leading cause of cancer deaths around the world (18). Approximately 85% of lung cancer are non-small cell lung cancer (NSCLC), with approximately 15% of lung cancer being small cell lung cancer (SCLC) (3). The main histological subtypes of NSCLC are adenocarcinoma and squamous cell carcinoma (3). If identified at an early stage, surgical resection of NSCLC offers a favorable prognosis, with 5-year survival rates of 70 - 90% for small, localized tumors (stage I) (5,16). However, approximately 75% patients have advanced disease at the time of diagnosis (stage III/IV) (20). As reported by the UK Office for National Statistics in 2014, patients diagnosed with distant metastatic disease (stage IV) had a 1-year survival rate of 15-19%. The majority of the patients of Adenocarcinomas - the most common subtype of lung cancer - come to clinical attention with distantly metastatic or locally advanced disease, since the disease may be asymptomatic in their early stages (17).

CT screening is a widely used diagnostic approach for lung cancer. However, in addition to the concerns of radiation exposures (1) and the cost of testing, CT screening is also associated with a high rate of falsely positive tests (1). Because early diagnosis is a key factor determining the 5-year survival rates of the disease, it becomes increasingly valuable to establish approaches that can conduct diagnosis of lung cancer at patients’ homes non-invasively and efficiently.

Skin autofluorescence (AF) has shown promise for non-invasive diagnosis of diabetes, which is based on detecting the AF of advanced glycation end-products (AGEs) of the collagen in dermis (10,13). The excitation wavelength of AGEs’ AF (9,11) is profoundly different from that of the epidermal AF of keratins (7,15). Keratins, together with melanin, NADH and FAD, are major epidermal fluorophores (2,15). Our recent study has found that UV-induced epidermal green AF, which is originated from UV-induced keratin 1 proteolysis in the spinous cells of epidermis, can be used as a novel biomarker for predicting UV-induced skin damage (7). Since detection of AF is non-invasive and simple, it is of great significance to further investigate the potential of AF as biomarkers for diseases. Our latest study has further found that the oxidative stress induced by UVC is causative to the increased epidermal AF of mouse ears by inducing keratin 1 proteolysis (12). Since a number of studies have found increased oxidative stress in the plasma of lung cancer patients (4,6,8,14,19), it is warranted to determine if there are increases in the epidermal AF of lung cancer patients, which may become a new diagnostic biomarker for the disease.

In this study, we tested our hypothesis that lung cancer patients may have increased AF in certain locations of their skin, which may become a novel diagnostic biomarker for the disease. Our study has provided first evidence that there are selective increases in green AF intensity at certain locations of the skin of lung cancer patients, which has provided evidence supporting our hypothesis.

## Methods

### Human subjects

This study was conducted according to a protocol approved by the Ethics Committee of Shanghai Chest Hospital, Shanghai Jiao Tong University, and a protocol approved by the Ethics Committee of Shanghai Fifth People’s Hospital, Fudan University. The human subjects in our study were divided into five groups: Group 1: The Healthy Group; Group 2: The Group of Pulmonary Infection Patients, who were hospitalized in the Department of Pulmonary Medicine, Shanghai Chest Hospital, Shanghai Jiao Tong University; Group 3: The Group of Untreated Lung Cancer Patients, including untreated adenocarcinoma, squamous-cell carcinoma and small-cell lung carcinoma patients, who were hospitalized in the Department of Pulmonary Medicine; Group 4: The Group of Chemotherapy-Treated Lung Cancer Patients who were hospitalized in the Department of Pulmonary Medicine; and Group 5: The Group of Acute Ischemic Stroke (AIS) patients who were diagnosed as AIS by the neurologists of the Department of Neurology, Shanghai Fifth People’s Hospital, Fudan University.

### Development of mouse model of lung cancer

Cell suspension of LLC (Lewis Lung Carcinoma) cells (1 × 10^6^ cells) in a total volume of 5 μl mixed with Matrigel (PBS: Matrigel = 4:1) were injected into the left lung of 4-week-old male C57BL/6 mice. One week after the injection, the mice were sacrificed and their lungs were obtained for determining if lung cancer was developed.

### Imaging of the AF of mouse’s skin

The AF of the ears of the mice were imaged by a two-photon fluorescence microscope (A1 plus, Nikon Instech Co., Ltd., Tokyo, Japan), with the excitation wavelength of 488 nm and the emission wavelength of 500 - 530 nm. The AF was quantified by the following approach: Sixteen spots with the size of approximately 100 × 100 μm^2^ on the scanned images were selected randomly. After the average AF intensities of each layer were calculated, the sum of the average AF intensities of all layers of each spot was calculated, which is defined as the AF intensity of each spot. If the value of average AF intensity of certain layer is below 45, the AF signal of the layer is deemed background noise, which is not counted into the sum.

### Determinations of the AF of human subjects

A portable AF imaging equipment was used to detect the AF of the fingernails and certain locations of the skin of the human subjects. The excitation wavelength is 485 nm, and the emission wavelength is 500 - 550 nm. For all of the human subjects, the AF intensity in the following seven locations on both hands, i.e., fourteen locations in total, was determined, including the index fingernails, Ventroforefingers, dorsal index fingers, Centremetacarpus, Dorsal Centremetacarpus, Ventriantebrachium, and Dorsal Antebrachium.

### Statistical analyses

All data are presented as mean + SEM. Data were assessed by one-way ANOVA, followed by Student - Newman - Keuls *post hoc* test, except where noted. *P* values less than 0.05 were considered statistically significant.

## Results

### 1. Development of lung cancer led to increased epidermal AF of mice

We studied the effects of development of lung cancer on the epidermal green AF of mice. We found that the with the development of lung cancer (Fig. 1), there was a marked increase in the epidermal green AF in all of the mice that had developed lung cancer (Fig. 2A). The approximately 3-fold increase in the AF intensity was statistically significant (Fig. 2B).

**Fig. 1.**
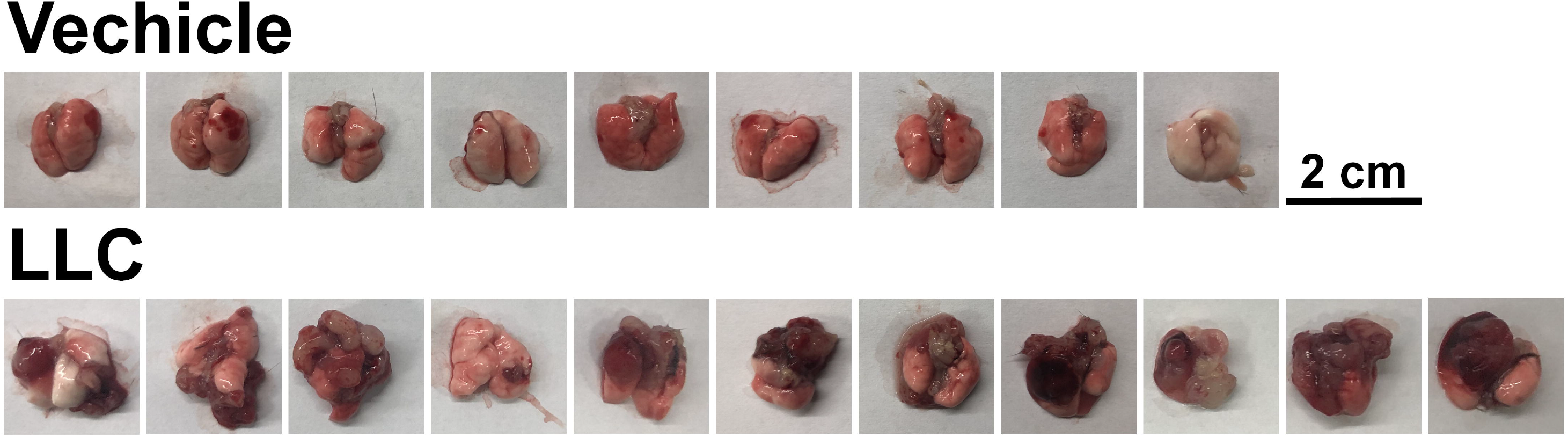
Development of orthotopic lung cancer led to increased epidermal AF of mice. One week after injection of the LLC cells into the lung of C57 mice, all of the mice developed lung cancer.

**Fig. 2.**
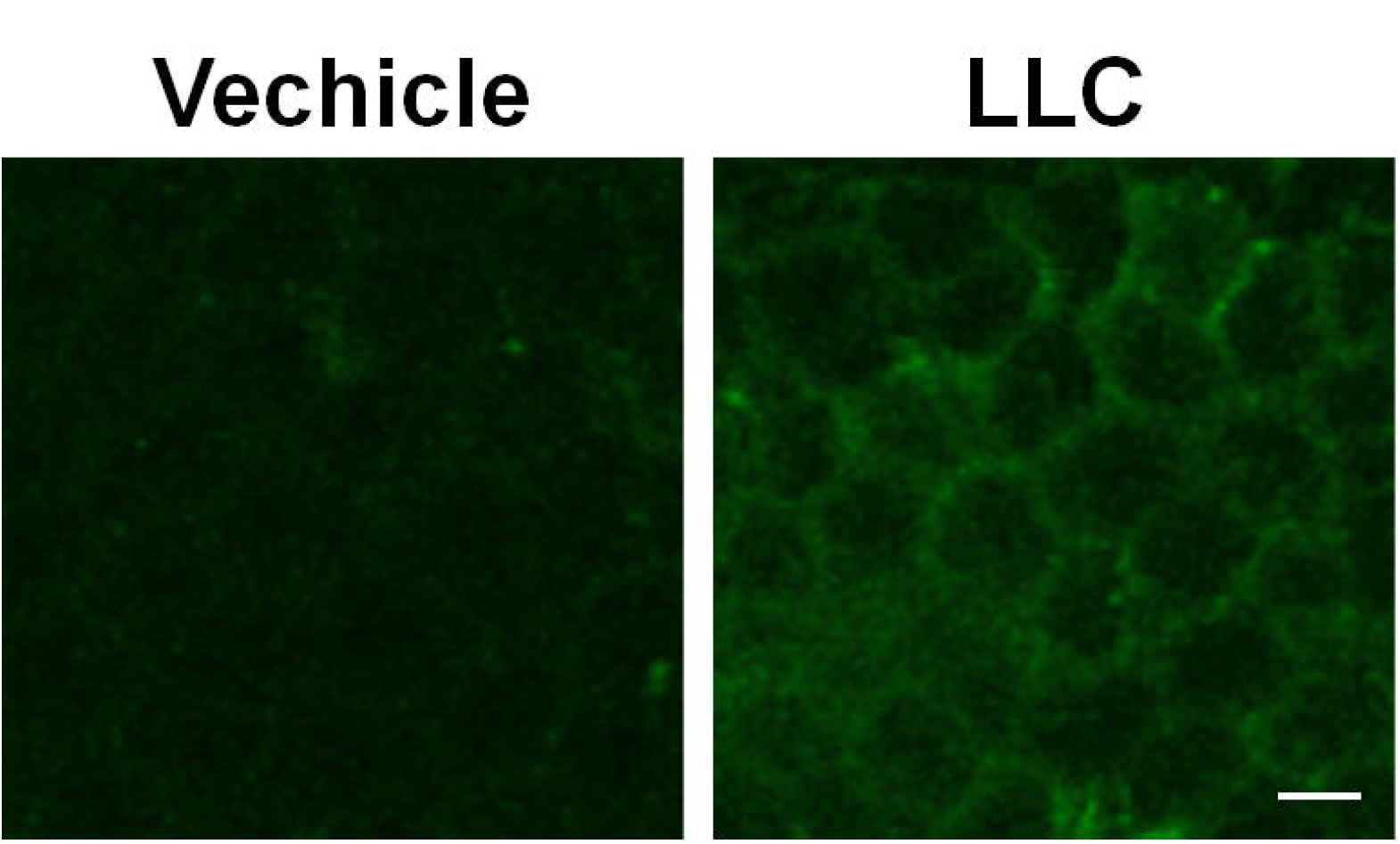
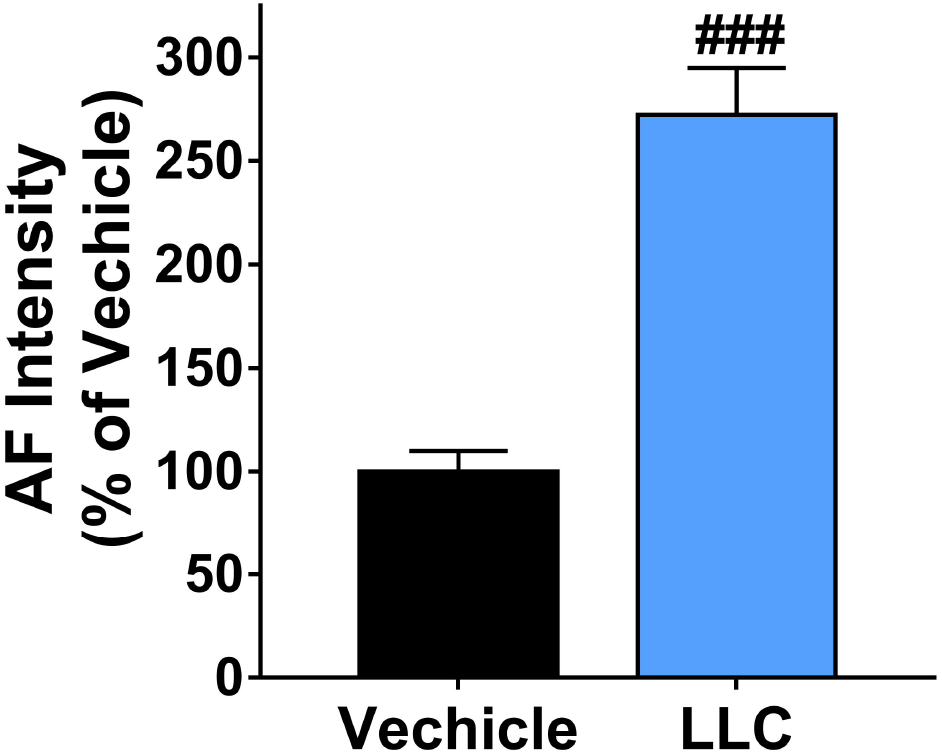
Development of lung cancer led to increased epidermal AF of mice. (A) There was a marked increase in the epidermal green AF in all of the mice that had developed lung cancer. (B) The approximately 3-fold increase in the AF intensity was statistically significant. N=10-12, ###, *P* < 0.001.

### 2. The AF intensity at certain locations of the skin of untreated lung cancer patients was significantly higher than that of the healthy persons and the patients of pulmonary infections

We determined the green AF intensity of two index fingernails and twelve locations of the skin of the Healthy Group, the Group of Pulmonary Infection Patients, and the Group of Untreated Lung Cancer Patients. We found that the AF intensity of the untreated lung cancer patients was significantly higher than that of the healthy persons at all of the examined positions (Fig. 3A – 3G). The AF intensity of the untreated lung cancer patients was also significantly higher than that of the pulmonary infection patients at right Ventriantebrachium (Fig. 3A), left and right Dorsal Antebrachium (Fig. 1B), and left and right Dorsal Centrimetacarpus (Fig. 3D).

**Fig. 3.**
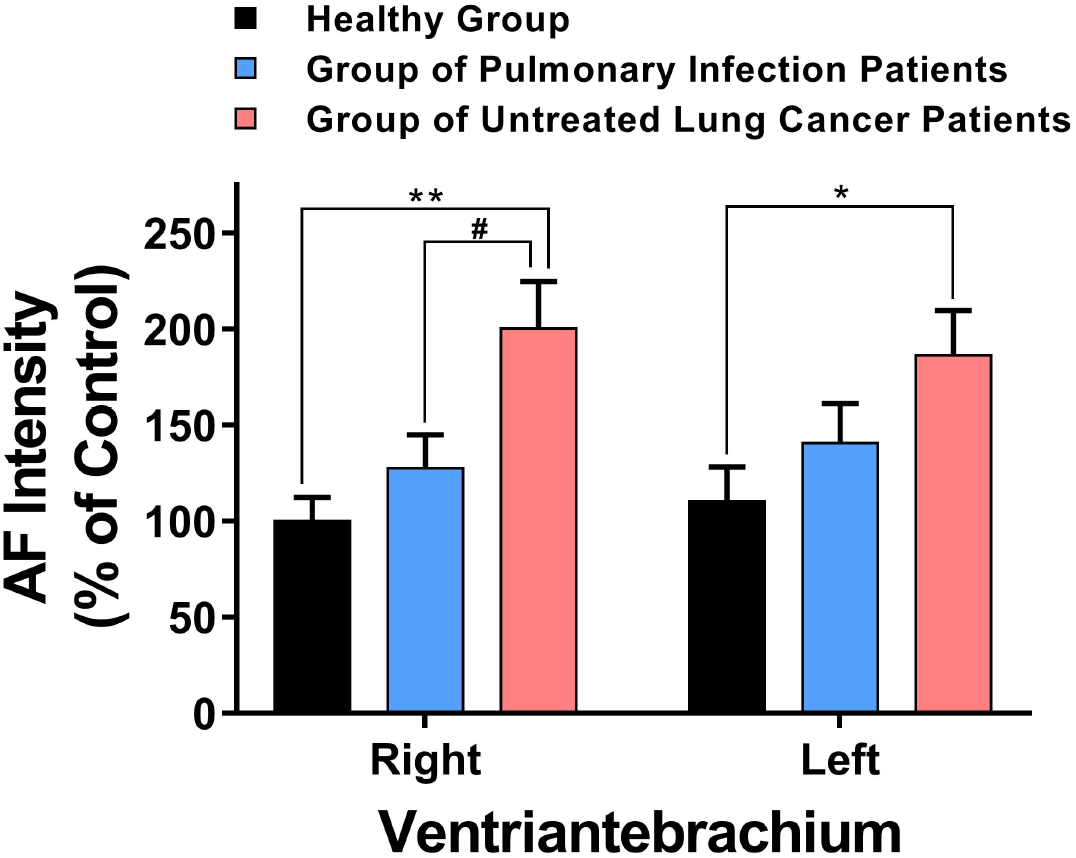
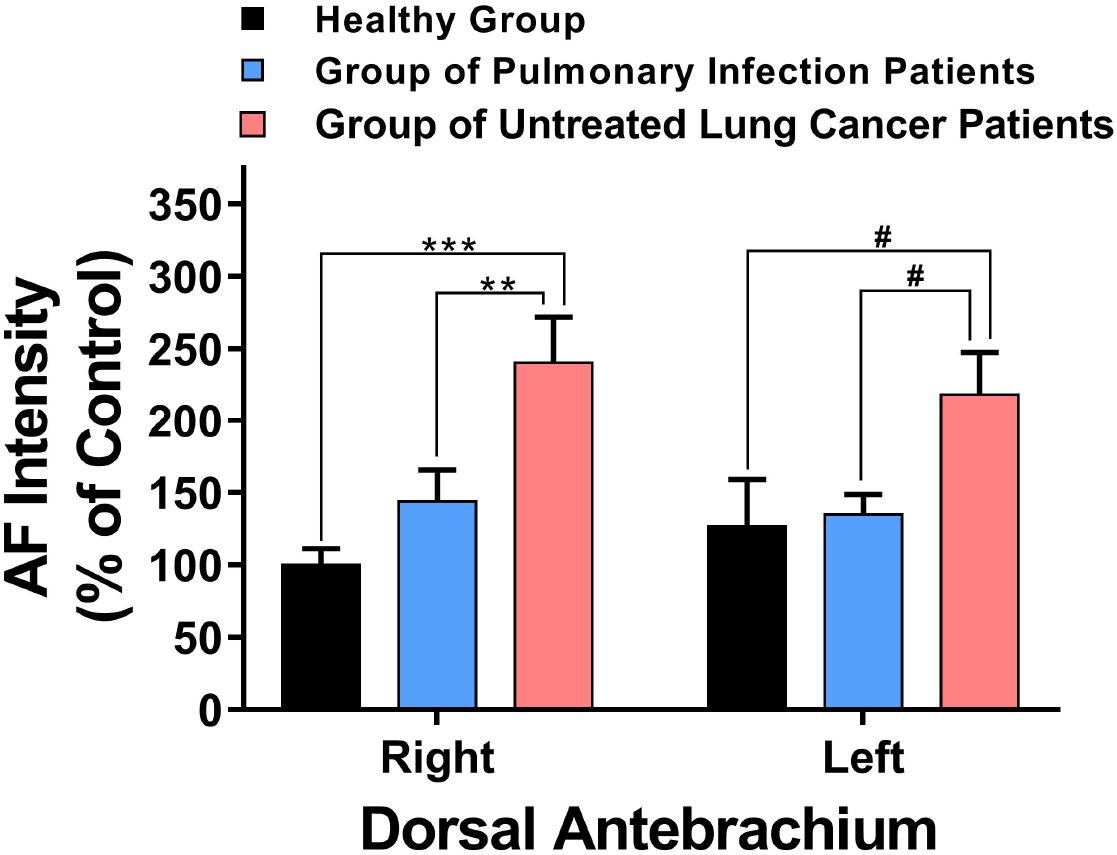
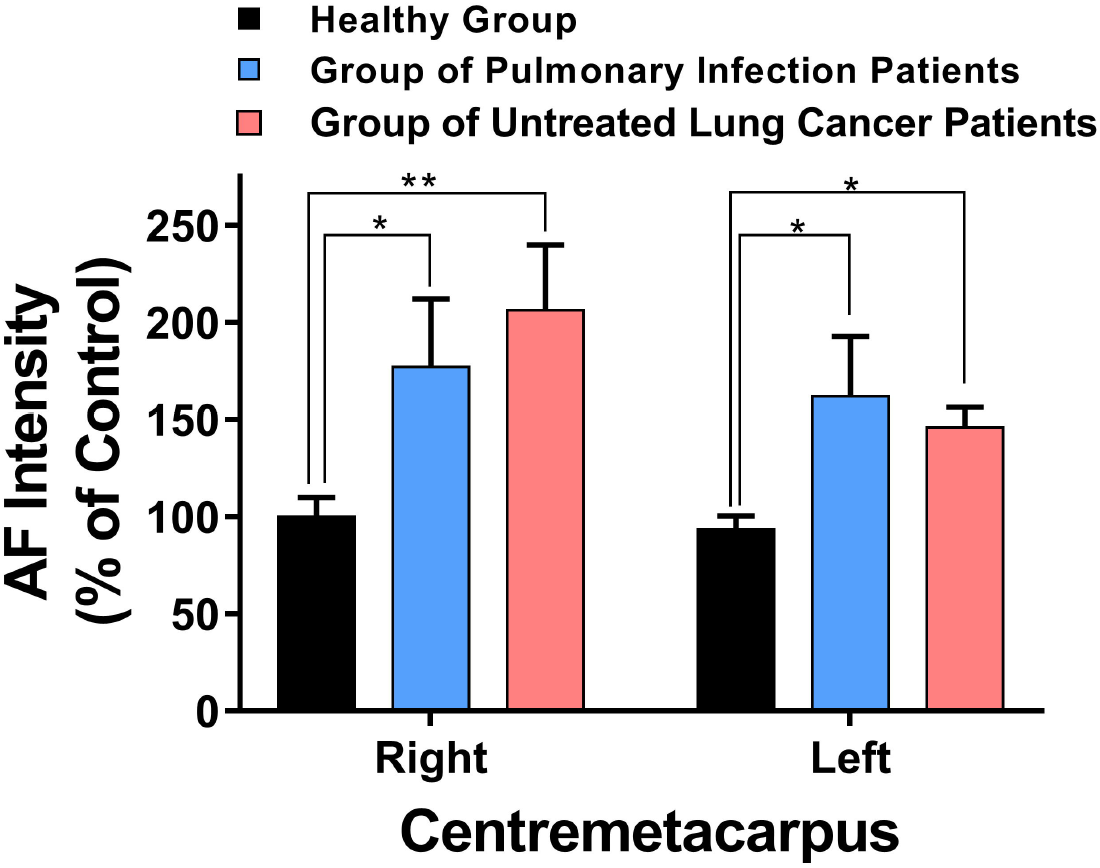
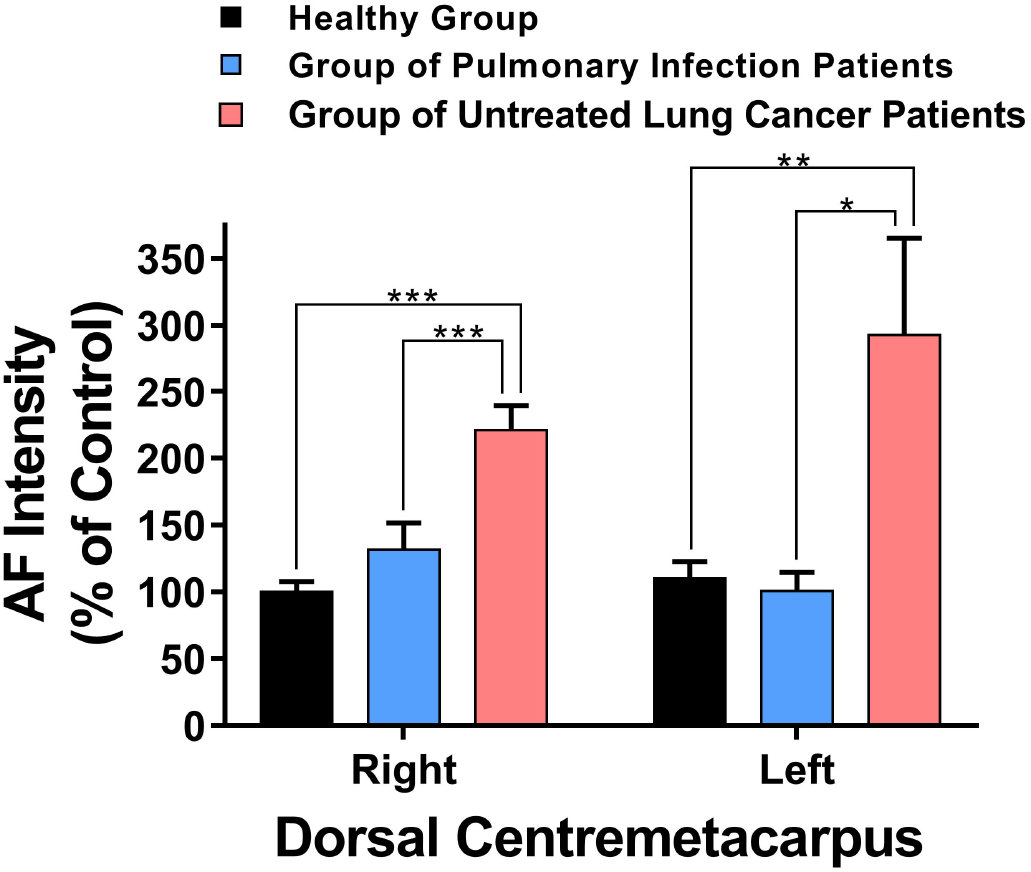
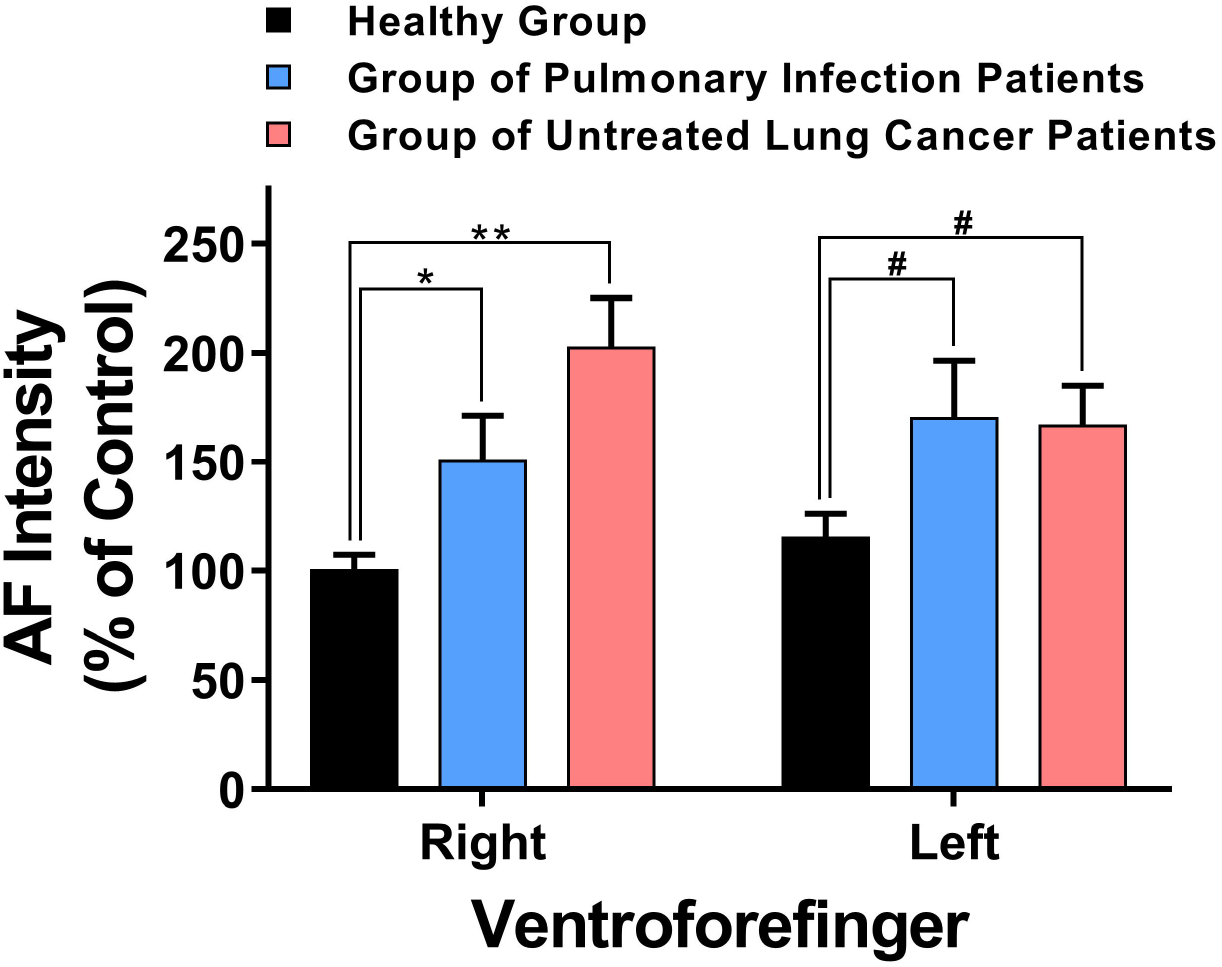
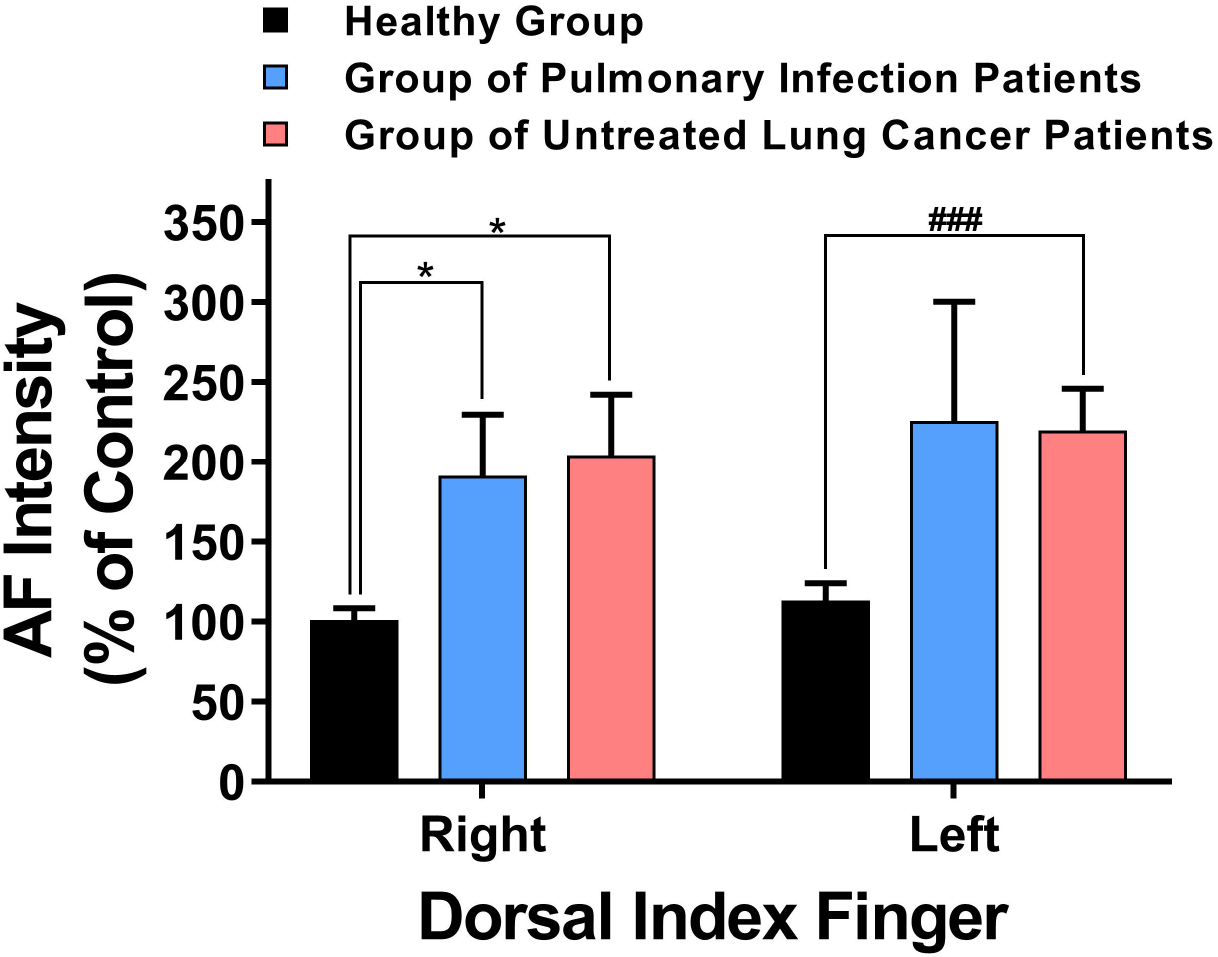
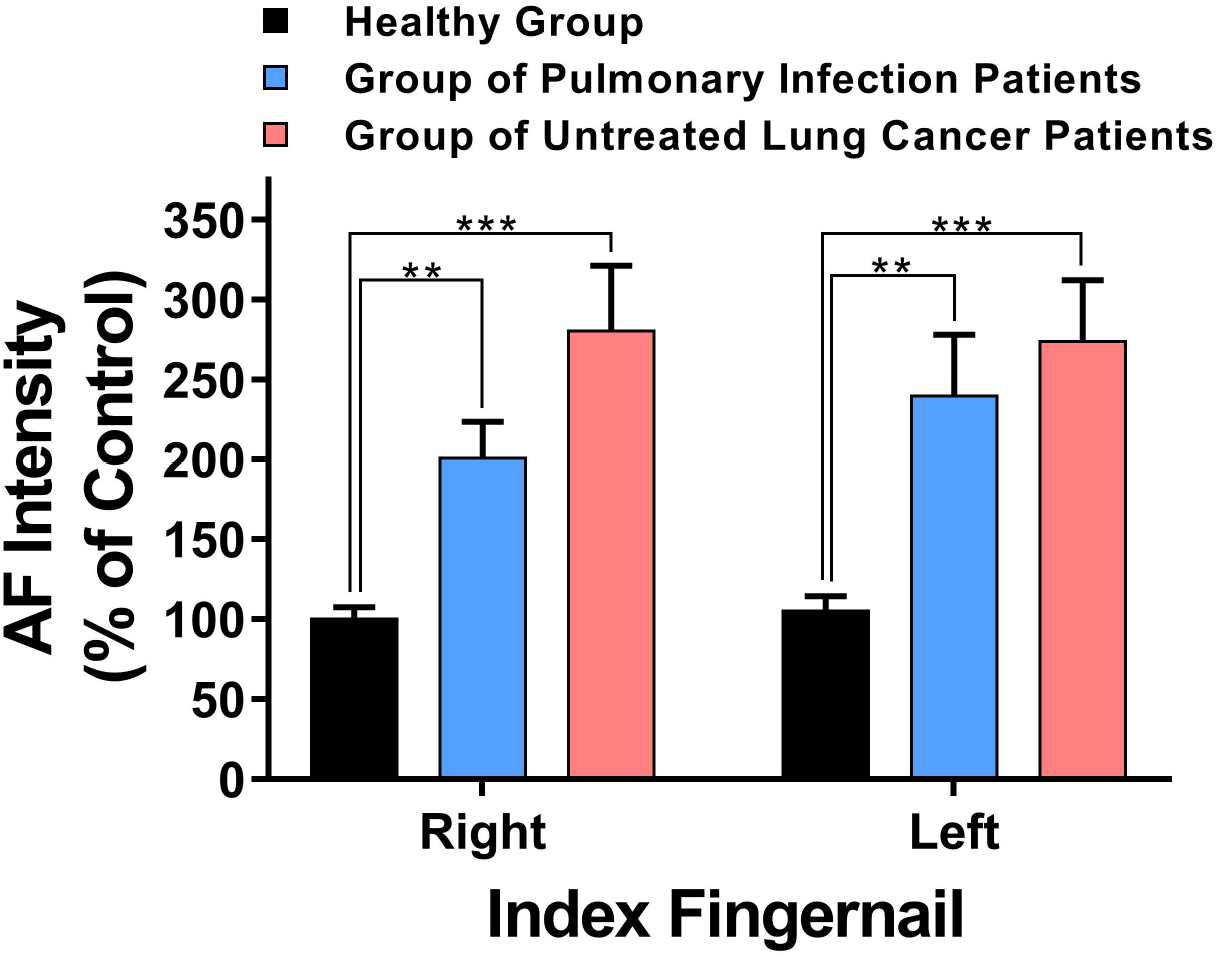
The AF intensity of certain examined locations of untreated lung cancer patients was significantly higher than that of the healthy persons and the pulmonary infection patients. (A) At right Ventriantebrachium, the AF intensity of the untreated lung cancer patients was significantly higher than that of the healthy persons and the pulmonary infection patients. At left Ventriantebrachium, the AF intensity of the untreated lung cancer patients was significantly higher than that of the healthy persons. (B) At both left and right Dorsal Antebrachium, the AF intensity of the untreated lung cancer patients was significantly higher than that of the healthy persons and the pulmonary infection patients. (C) At both left and right Centrimetacarpus, the AF intensity of the untreated lung cancer patients was significantly higher than that of the healthy persons. (D) At both left and right Dorsal Centrimetacarpus, the AF intensity of the untreated lung cancer patients was significantly higher than that of the healthy persons and the pulmonary infection patients. (E) At both left and right Ventroforefinger, the AF intensity of the untreated lung cancer patients was significantly higher than that of the healthy persons. (F) At both left and right Dorsal Index Finger, the AF intensity of the untreated lung cancer patients was significantly higher than that of the healthy persons. (G) At both left and right Index Fingernails, the AF intensity of the untreated lung cancer patients was significantly higher than that of the healthy persons. The number of subjects in the Healthy Group, the Group of Pulmonary Infection Patients, and the Group of Untreated Lung Cancer Patients was 54, 29, and 39, respectively. *,*p* < 0.05; **,*p* < 0.01; ***, *p* < 0.001; #, *p* < 0.05 (Student *t*-test); ##, *p* < 0.01 (Student *t*-test); ###, *p* < 0.001 (Student *t*-test).

### 3. The AF asymmetry of untreated lung cancer patients was significantly different from that of the healthy persons and the pulmonary infection patients at certain examined locations

We determined the ‘asymmetry of the AF intensity’, which is defined as ‘ difference between the AF intensity of the symmetrical positions of the body’. There was no significant difference among the AF asymmetry of the Healthy Group, the Group of Pulmonary Infection Patients, and the Group of Untreated Lung Cancer Patients at Ventriantebrachium (Fig. 4A), Dorsal Antebrachium (Fig. 4B) and Dorsal Centrimetacarpus (Fig. 4D). The AF asymmetry of both the Group of Pulmonary Infection Patients and the Group of Untreated Lung Cancer Patients was significantly higher than that of the Healthy Group at Centrimetacarpus (Fig. 4C) and Index Fingernail (Fig. 4G). The AF asymmetry of the Group of Pulmonary Infection Patients was significantly higher than that of the Healthy Group at Ventroforefinger (Fig. 4E). At Dorsal Index Fingers, the AF asymmetry of the Group of Pulmonary Infection Patients was significantly higher than that of the Healthy Group and the Group of Untreated Lung Cancer Patients (Fig. 4F).

**Fig. 4.**
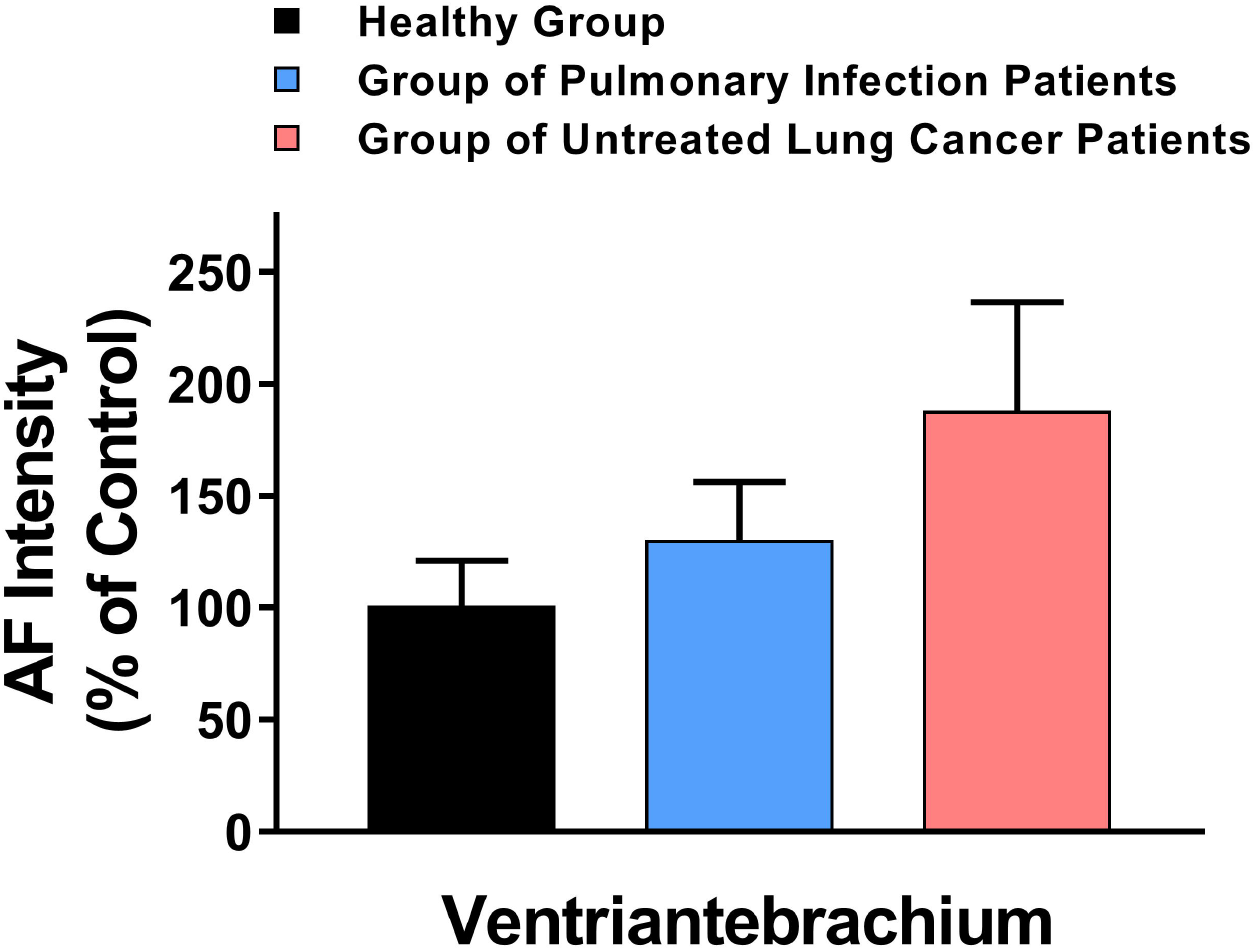
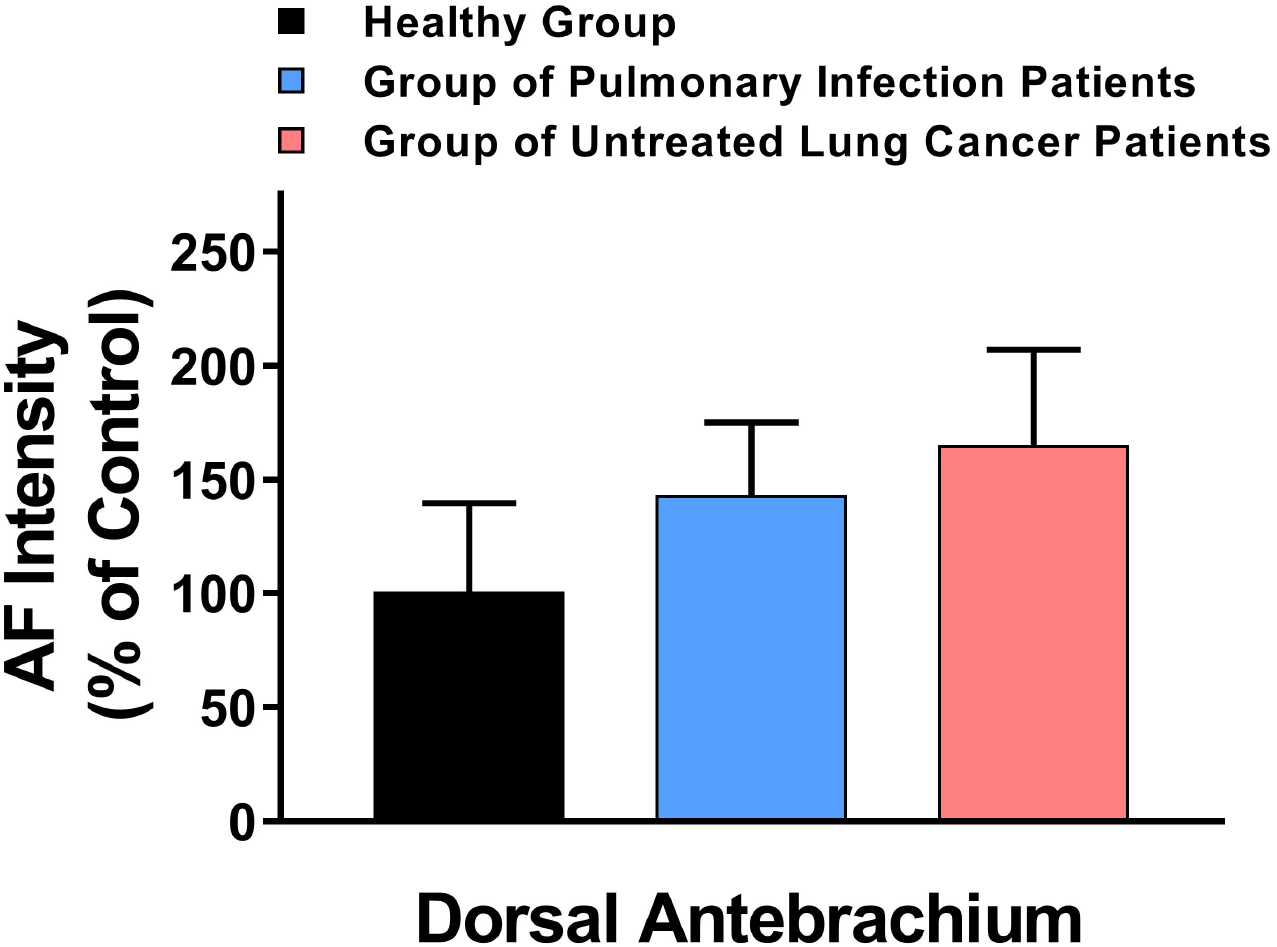
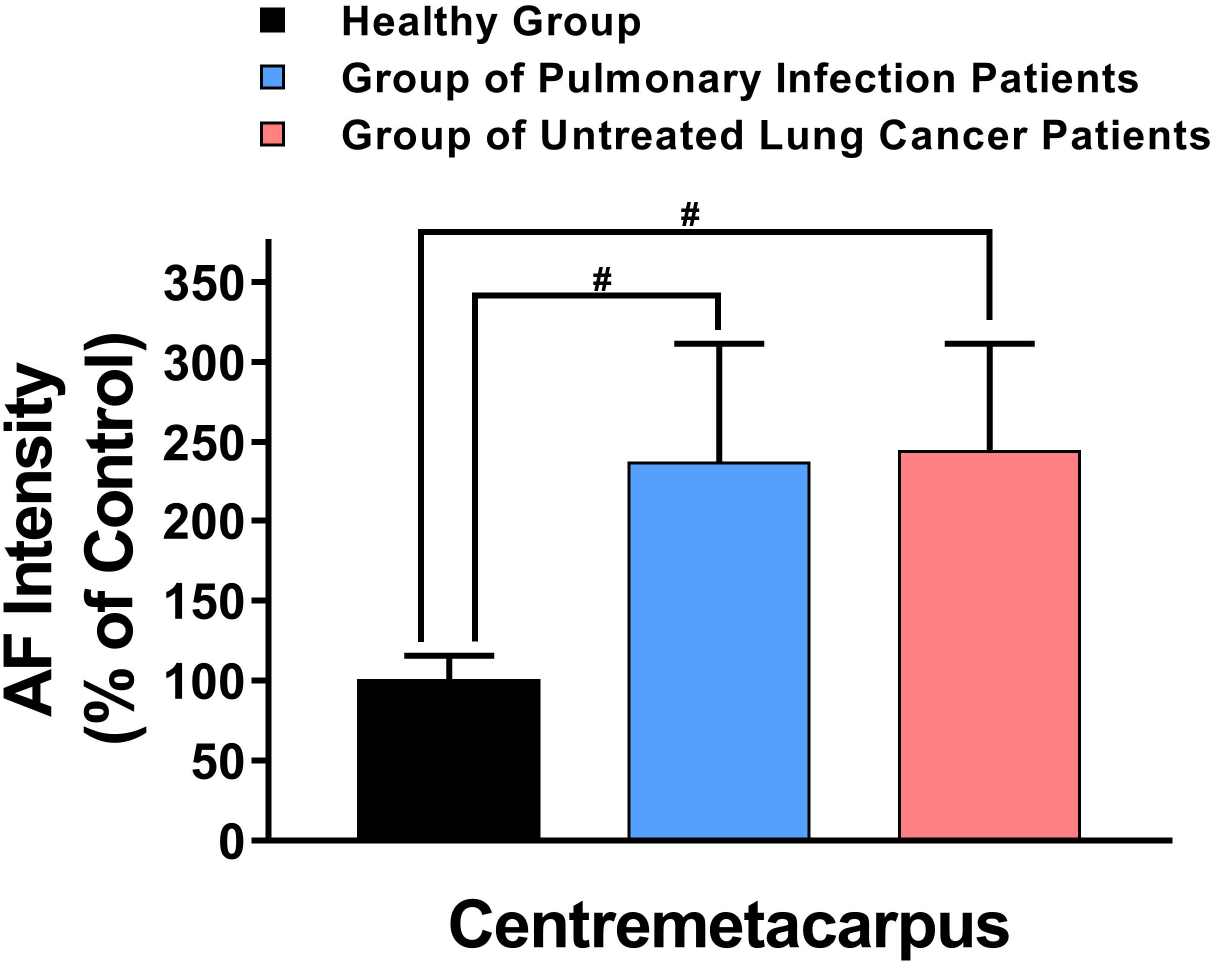
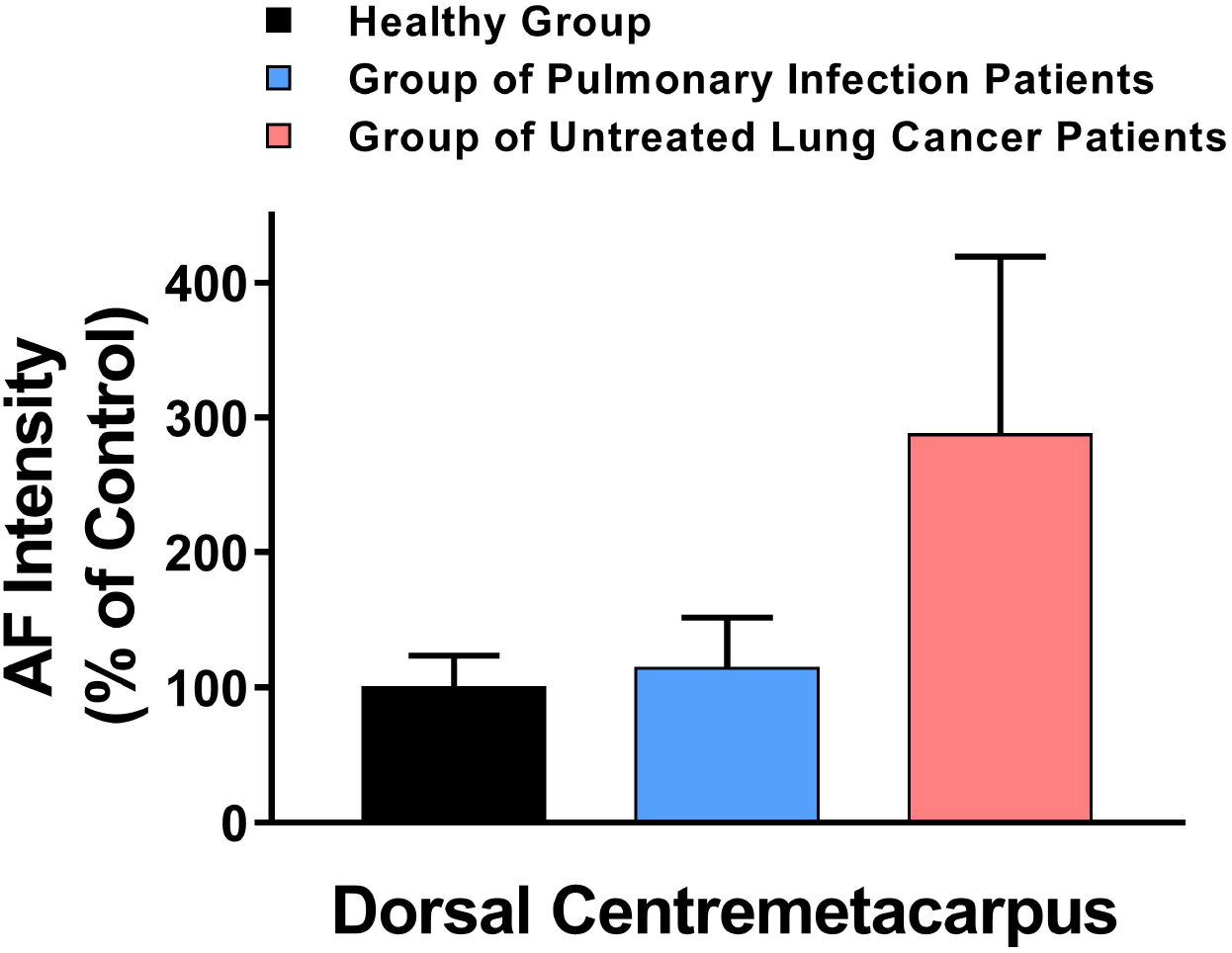
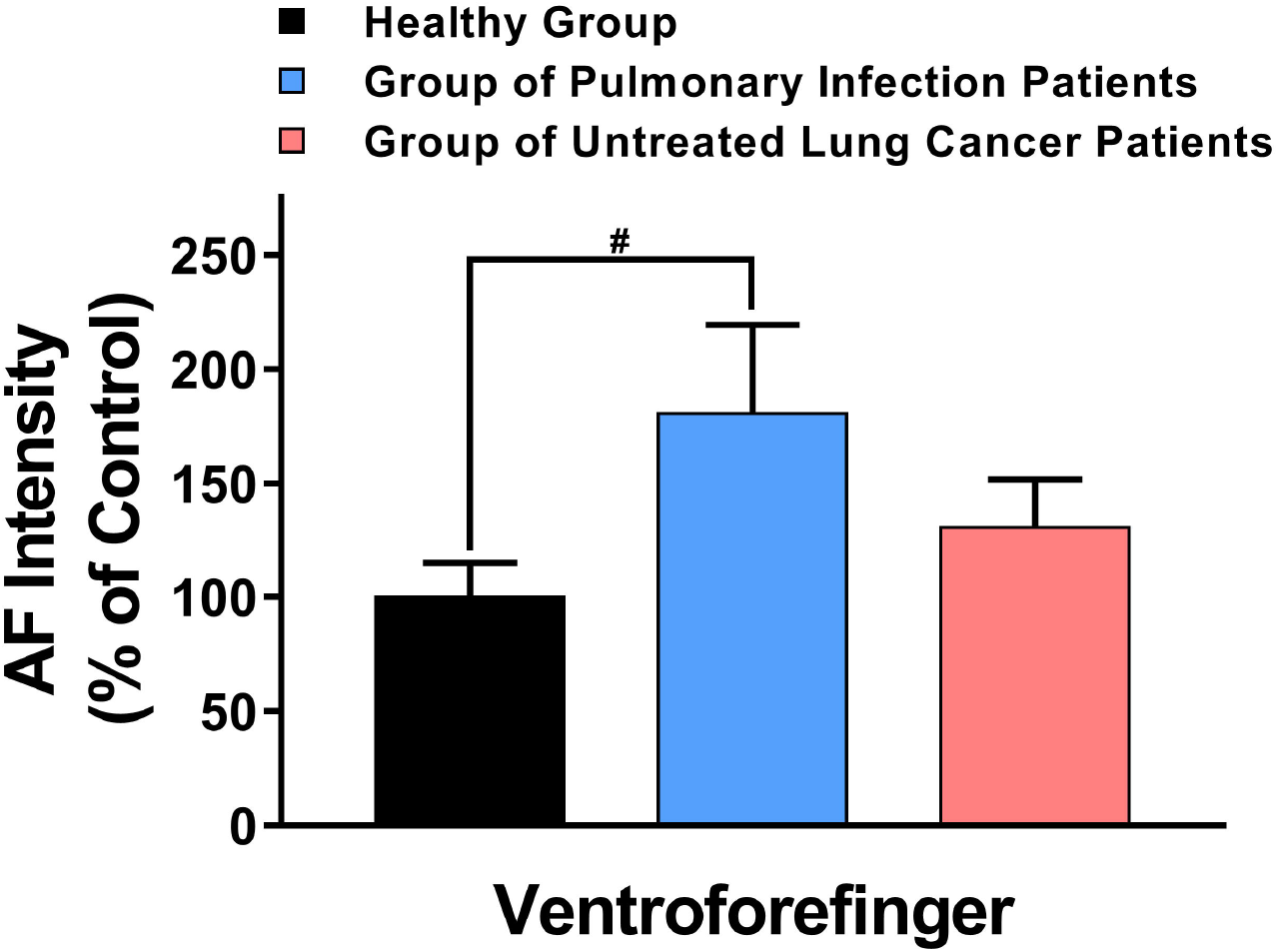
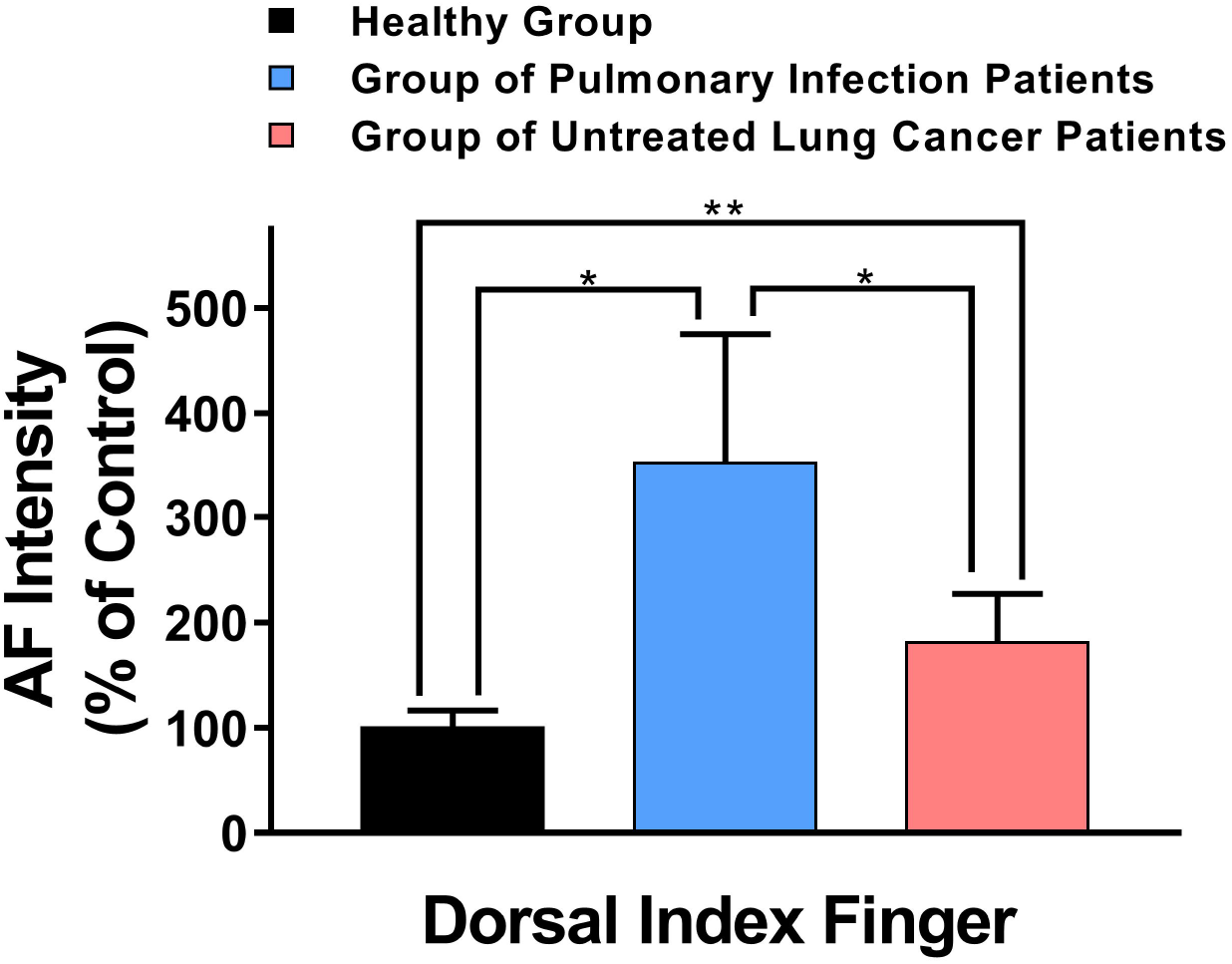
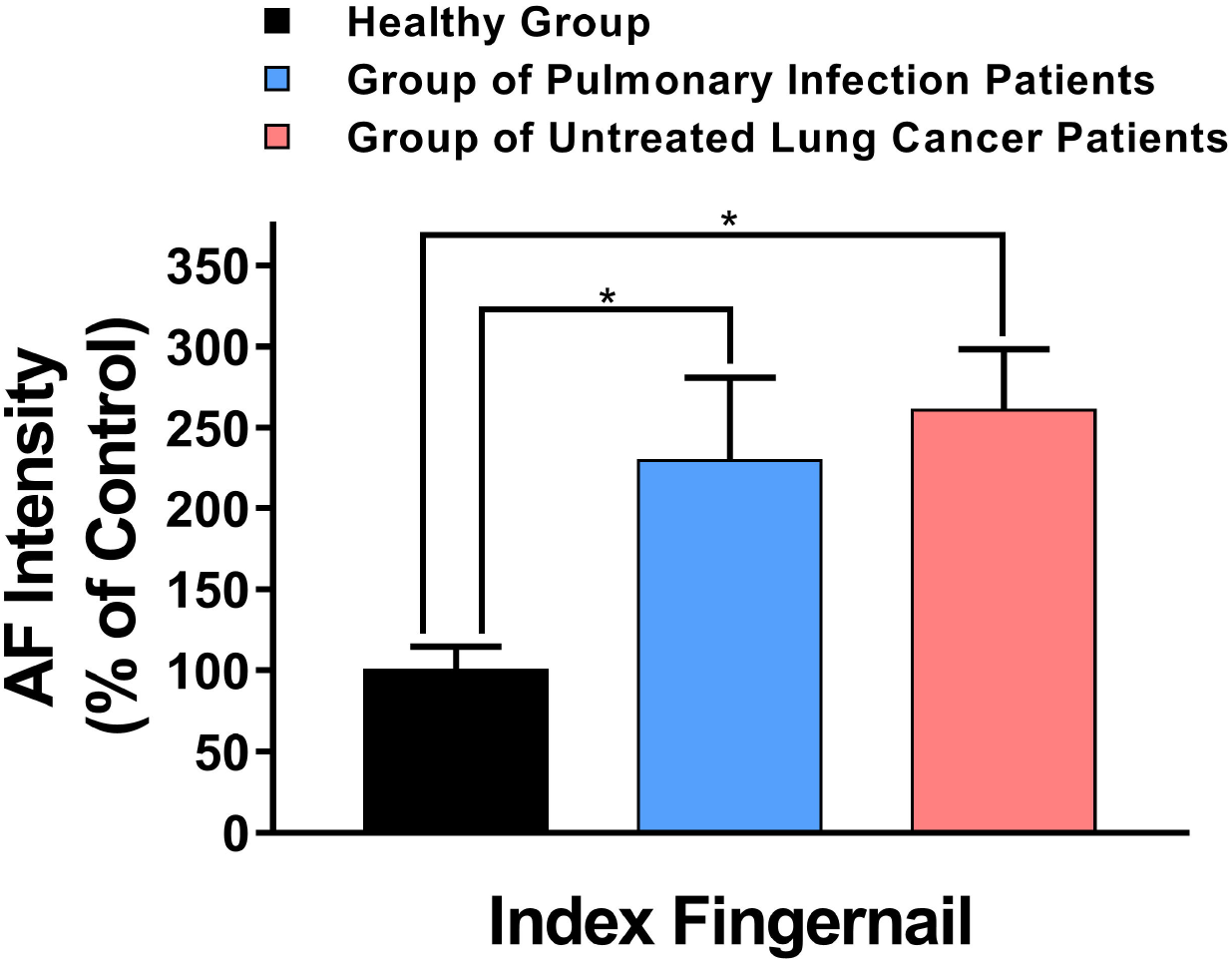
The AF asymmetry of untreated lung cancer patients was significantly different from that of the healthy persons and the pulmonary infection patients at certain examined locations. There was no significant difference among the AF asymmetry of the Healthy Group, the Group of Pulmonary Infection Patients, and the Group of Untreated Lung Cancer Patients at Ventriantebrachium (A), Dorsal Antebrachium (B) and Dorsal Centrimetacarpus (D). The AF asymmetry of both the Group of Pulmonary Infection Patients and the Group of Untreated Lung Cancer Patients was significantly higher than that of the Healthy Group at Centrimetacarpus (C) and Index Fingernail (G). The AF asymmetry of the Group of Pulmonary Infection Patients was significantly higher than that of the Healthy Group at Ventroforefinger (E). At Dorsal Index Fingers, the AF asymmetry of the Group of Pulmonary Infection Patients was significantly higher than that of the Healthy Group and the Group of Untreated Lung Cancer Patients (F). The number of subjects in the Healthy Group, the Group of Pulmonary Infection Patients, and the Group of Untreated Lung Cancer Patients was 54, 29, and 39, respectively. *,*p* < 0.05; **,*p* < 0.01; ***,*p* < 0.001;#, *p* < 0.05 (Student *t*-test).

### 4. ‘The pattern of AF’ of the untreated lung cancer was distinctly different from that of the healthy persons and the pulmonary infection patients

We found that the percentage of the people with no more than one region with increased green AF was 50.0%, 24.1% and 2.6% for the Healthy Group, the Group of Pulmonary Infection Patients, and the Group of Untreated Lung Cancer Patients, respectively (Fig. 5A). In contrast, the percentage of the people with more than six locations with increased green AF was 7.4%, 20.7% and 61.5% for the Healthy Group, the Group of Pulmonary Infection Patients, and the Group of Untreated Lung Cancer Patients, respectively (Fig. 5A).

**Fig. 5.**
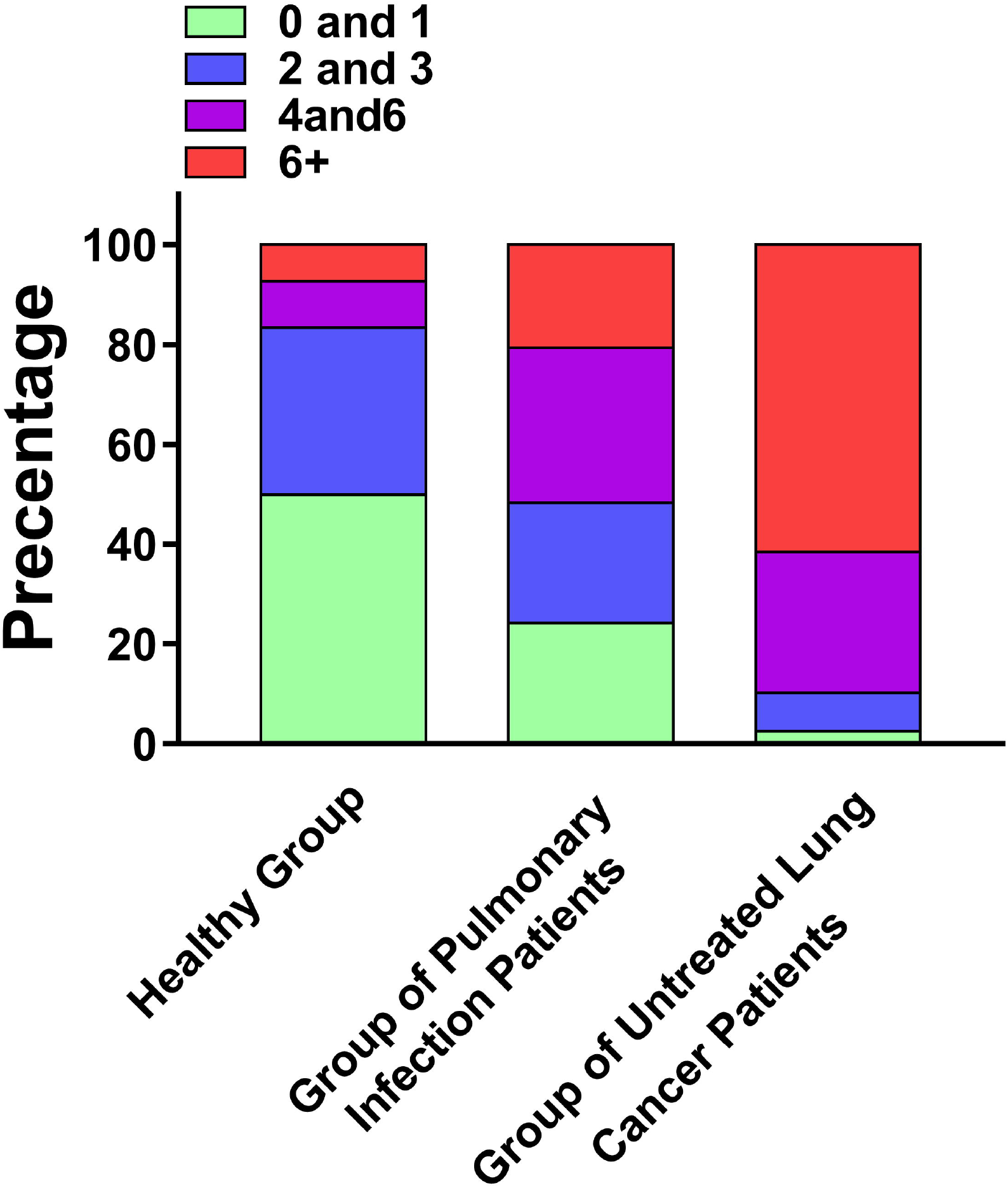
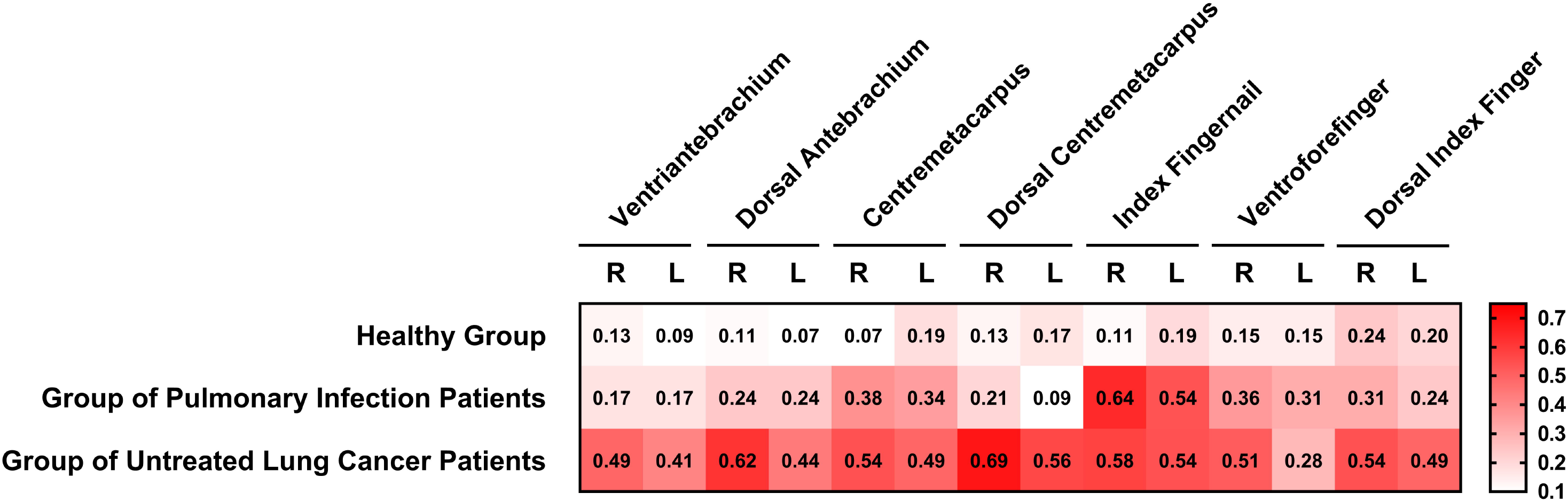
Comparisons of certain properties of the AF of the Healthy Group, the Group of Pulmonary Infection Patients, and the Group of Untreated Lung Cancer Patients. (A) The percentages of the people with 0-1, 2-3, 4-6, or 6+ locations with increased AF intensity were markedly different among the Healthy Group, the Group of Pulmonary Infection Patients, and the Group of Untreated Lung Cancer Patients. (B) In all of the locations examined, a markedly higher percentage of the untreated lung cancer patients had increased AF intensity compared with the healthy persons. In a majority of the locations examined, a higher percentage of the untreated lung cancer patients had increased AF intensity compared with the pulmonary infection patients. In the scale of the figure, 0.1, 0.2, 0.3, 0.4, 0.5 denotes 10%, 20%, 30%, 40%, 50%. The number of subjects in the Healthy Group, the Group of Pulmonary Infection Patients, and the Group of Untreated Lung Cancer Patients was 54, 29, and 39, respectively.

In all of the locations examined, a markedly higher percentage of the untreated lung cancer patients had increased AF intensity compared with the healthy persons (Fig. 5B). Moreover, in a majority of the locations examined, a higher percentage of the untreated lung cancer patients had increased AF intensity compared with the pulmonary infection patients (Fig. 5B).

As a major index of ‘Pattern of AF’, the green AF intensity of all of the examined locations of the three studied groups was listed in Table 1. The table showed that the ‘Pattern of AF’ was markedly different among the three groups. Moreover, the ‘Pattern of AF’ of the untreated lung cancer patients was significantly different from that of AIS patients (Table 2).

**Table 1.** The ‘Pattern of AF’ was markedly different among the the Healthy Group, the Group of Pulmonary Infection Patients, and the Group of Untreated Lung Cancer Patients. The number of subjects in the Healthy Group, the Group of Pulmonary Infection Patients, and the Group of Untreated Lung Cancer Patients was 54, 29, and 39, respectively.

| Factors |  | Healthy Group<br>(n=54) | Pulmonary Infection<br>Group (n=29) | Untreated Lung<br>Cancer Group<br>(n=39) |
| --- | --- | --- | --- | --- |
| AF of Ventroforefinger | R | 9.12±0.67 | 13.72±1.9<sup>##</sup> | 18.46±2.09<sup>###</sup> |
|  | L | 10.49±1.02 | 15.5±2.44<sup>#</sup> | 15.19±1.7<sup>#</sup> |
| AF of Index Fingernail | R | 6.28±0.49 | 12.61±1.44<sup>###</sup> | 17.61±2.56<sup>###</sup> |
|  | L | 6.6±0.58 | 15.05±2.41<sup>###</sup> | 17.2±2.4<sup>###</sup> |
| AF of Dorsal Index Finger | R | 5.28±0.45 | 10.06±2.08<sup>##</sup> | 10.73±2.07<sup>##</sup> |
|  | L | 5.93±0.62 | 11.87±3.99 | 11.55±1.44<sup>###</sup> |
| AF of Ventriantebrachium | R | 6±0.74 | 7.64±1.06 | 12±1.47<sup>###</sup> |
|  | L | 6.61±1.07 | 8.44±1.23 | 11.17±1.39<sup>##</sup> |
| AF of Dorsal Antebrachium | R | 4.64±0.53 | 6.68±1.01 | 11.16±1.46<sup>###</sup> |
|  | L | 5.88±1.5 | 6.26±0.66 | 10.12±1.37<sup>#</sup> |
| AF of Centremetacarpus | R | 6.15±0.6 | 10.89±2.16<sup>##</sup> | 12.67±2.08<sup>###</sup> |
|  | L | 5.75±0.43 | 9.95±1.91<sup>##</sup> | 8.97±0.65<sup>###</sup> |
| AF of Dorsal Centremetacarpus | R | 3.54±0.28 | 4.66±0.72 | 7.82±0.65<sup>###</sup> |
|  | L | 3.89±0.46 | 3.57±0.5 | 10.37±2.57<sup>##</sup> |

AF Intensity of the healthy Group, the pulmonary infection group and untreated lung cancer group. (student t-test, two tail)

**Table 2.**
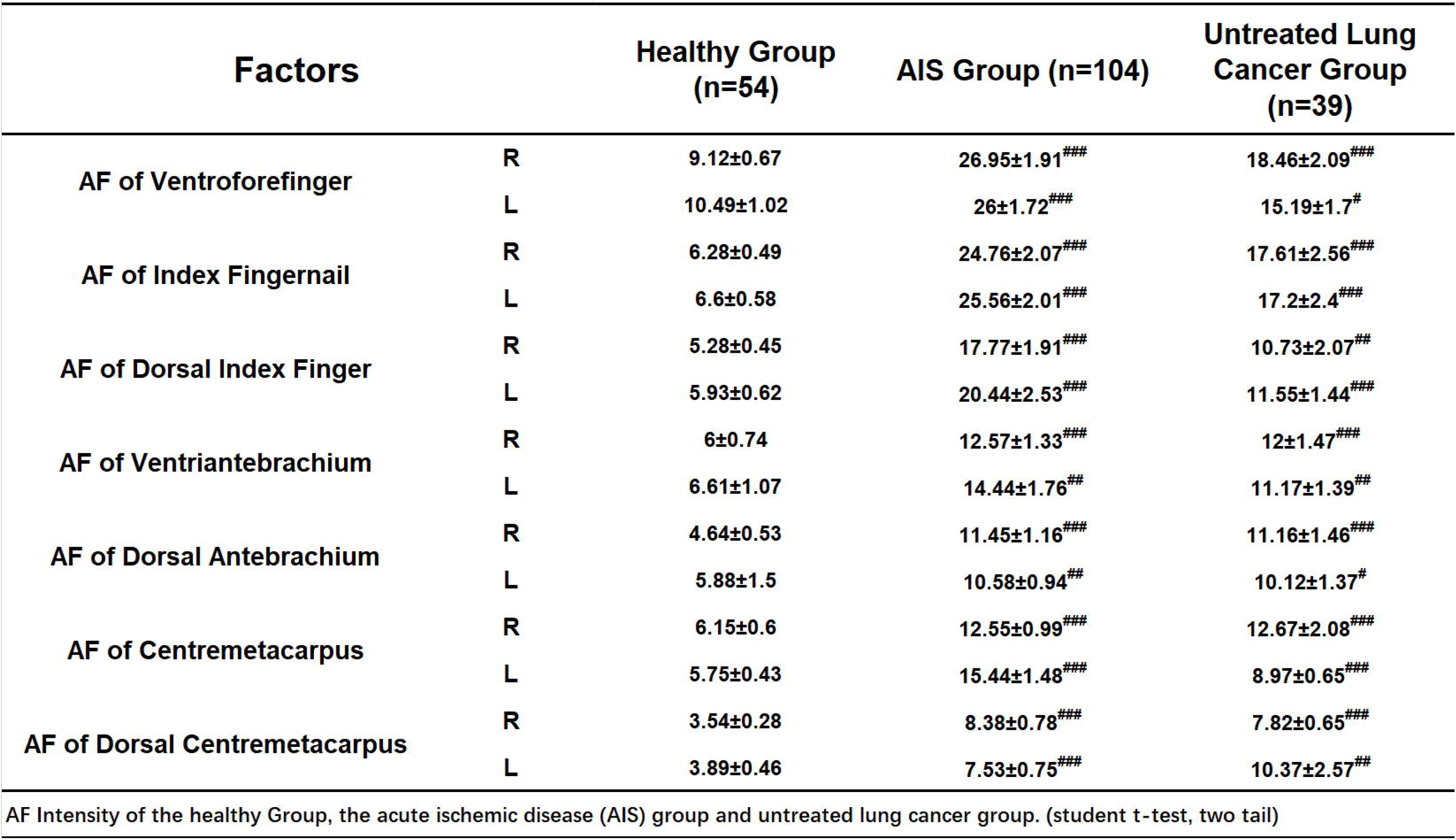
The ‘Pattern of AF’ of the untreated lung cancer patients was significantly different from that of acute ischemic stroke (AIS) patients. The number of subjects in the Healthy Group, the AIS Group, and the Group of Untreated Lung Cancer Patients was 54, 104, and 39, respectively.

### 4. ROC analyses indicated significant promise of the green AF-based technology in differentiating untreated lung cancer patients from either healthy persons or pulmonary infection patients

By using the AF intensity of each examined location as the sole diagnostic parameter, we conducted ROC analyses to determine the AUC for differentiating the healthy persons and untreated lung cancer patients. The AUC ranged from 0.8196 to 0.878 when the AF intensity of anyone of the examined locations, except right and left Ventroforefinger and left Centremetacarpus, was used as the sole diagnostic parameter (Fig. 6A – 6G). We also conducted ROC analyses to determine the AUC for differentiating the pulmonary infection patients and untreated lung cancer patients. The AUC was 0.8046 and 0.8841, respectively, when the AF intensity of right or left Dorsal Centremetacarpus was used as the sole diagnostic parameter (Fig. 6D).

**Fig. 6.**
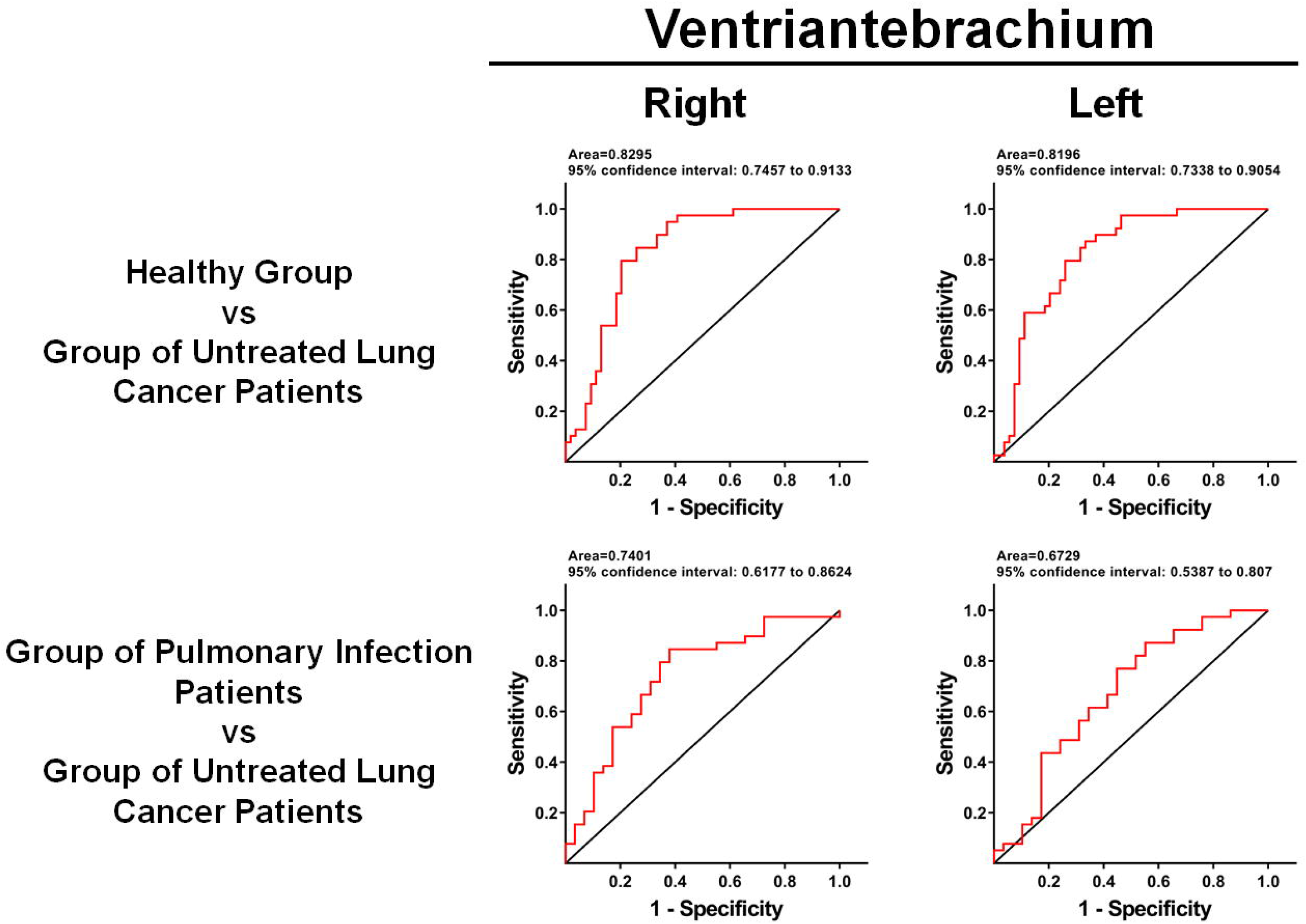
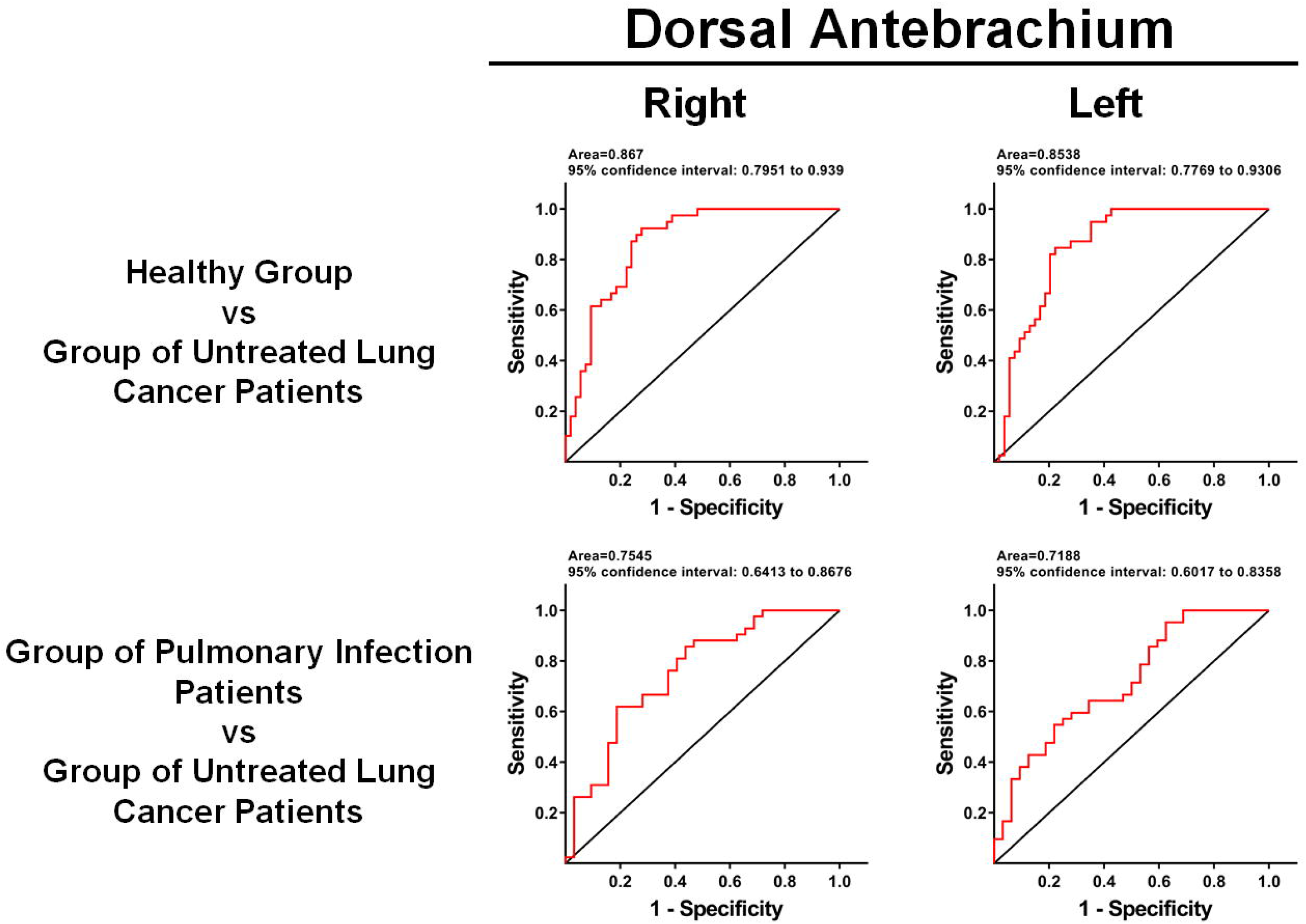
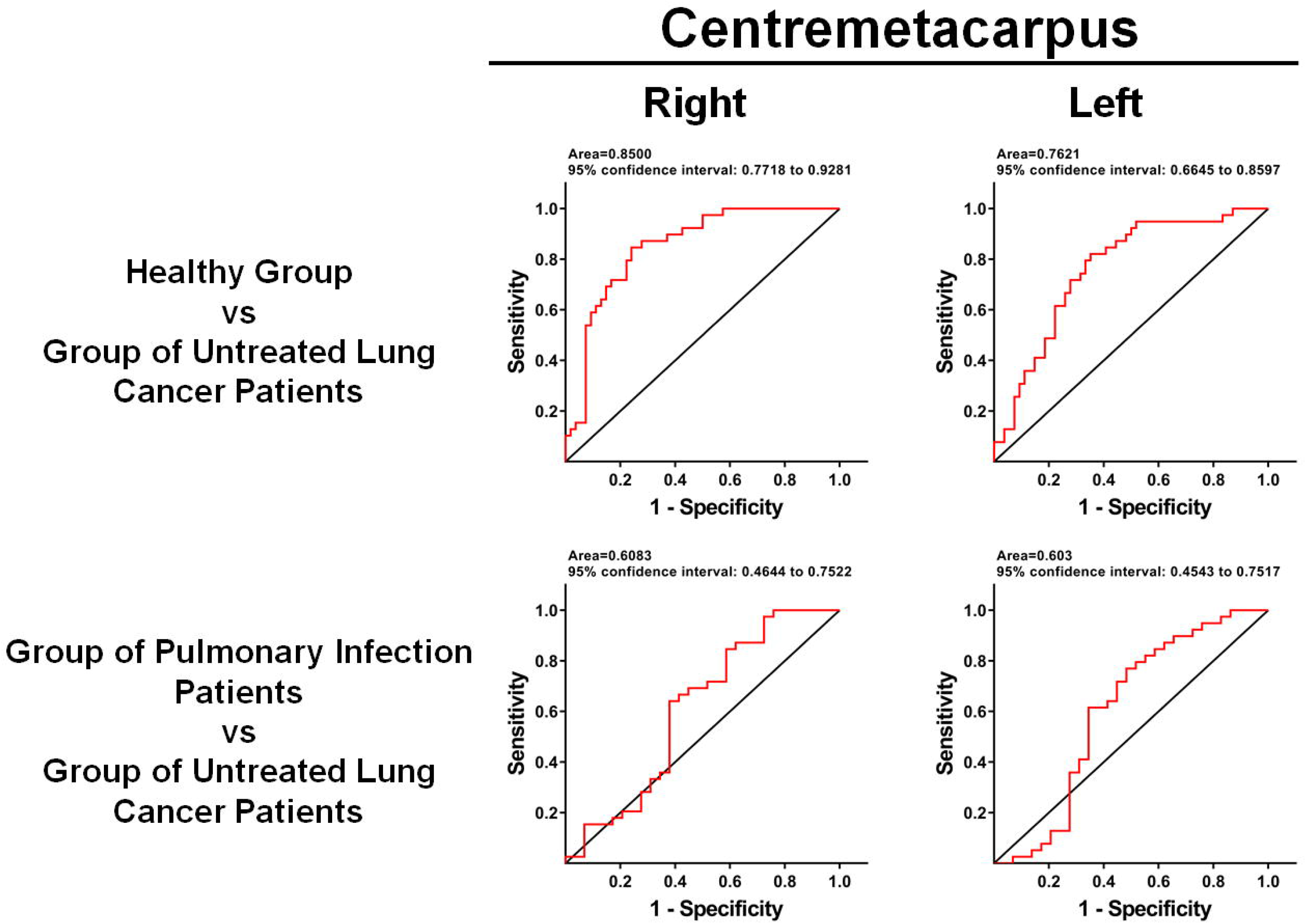
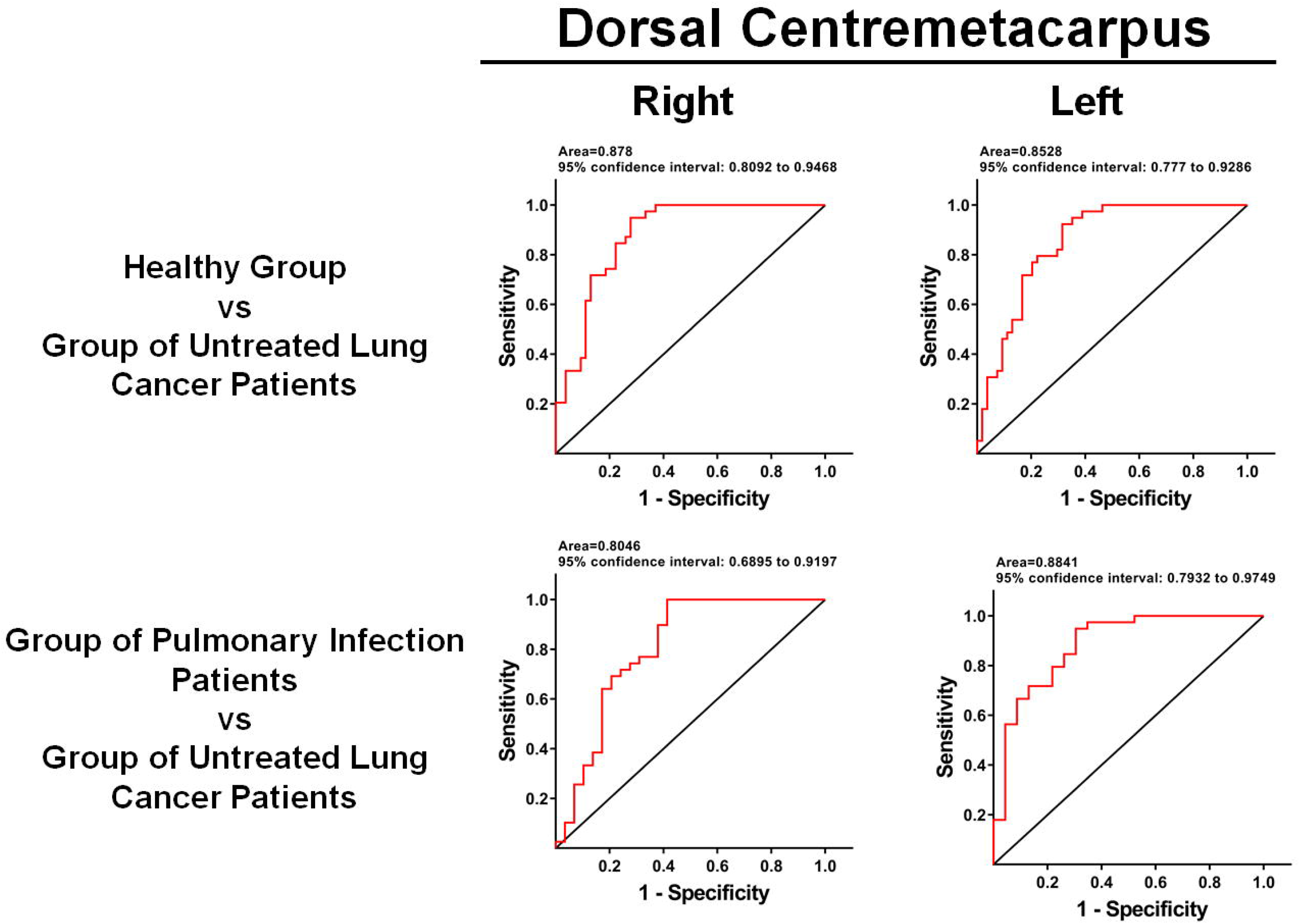
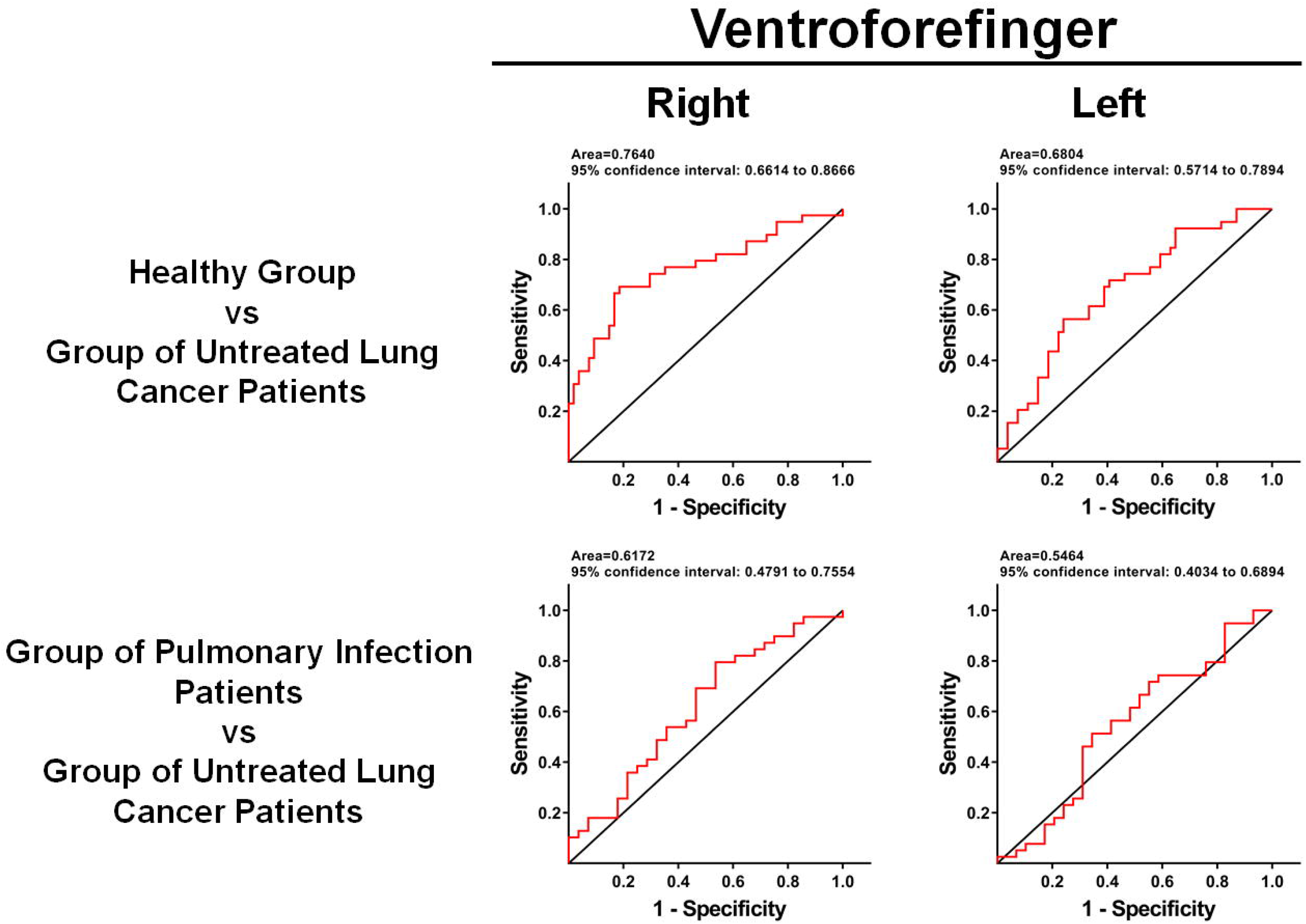
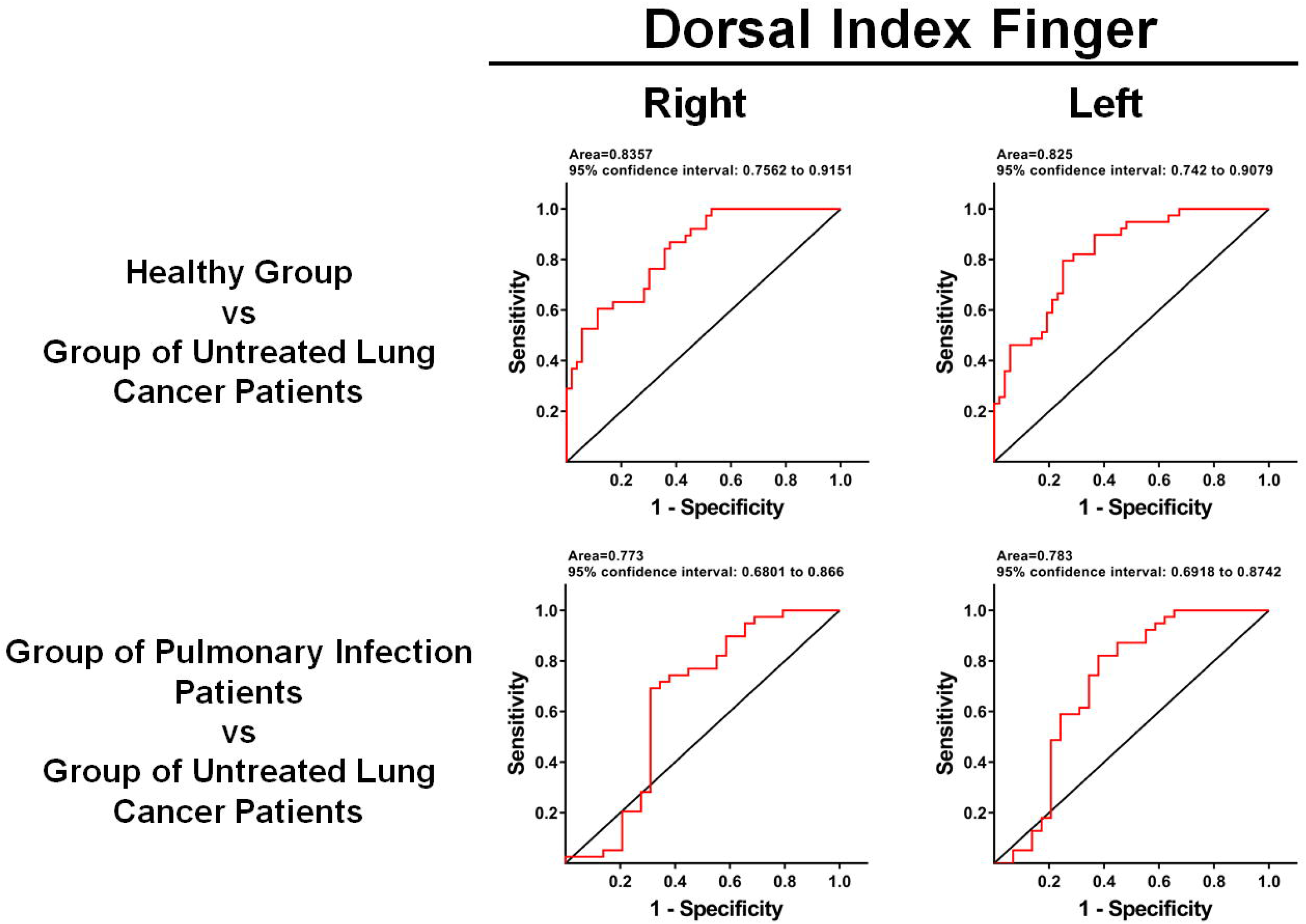
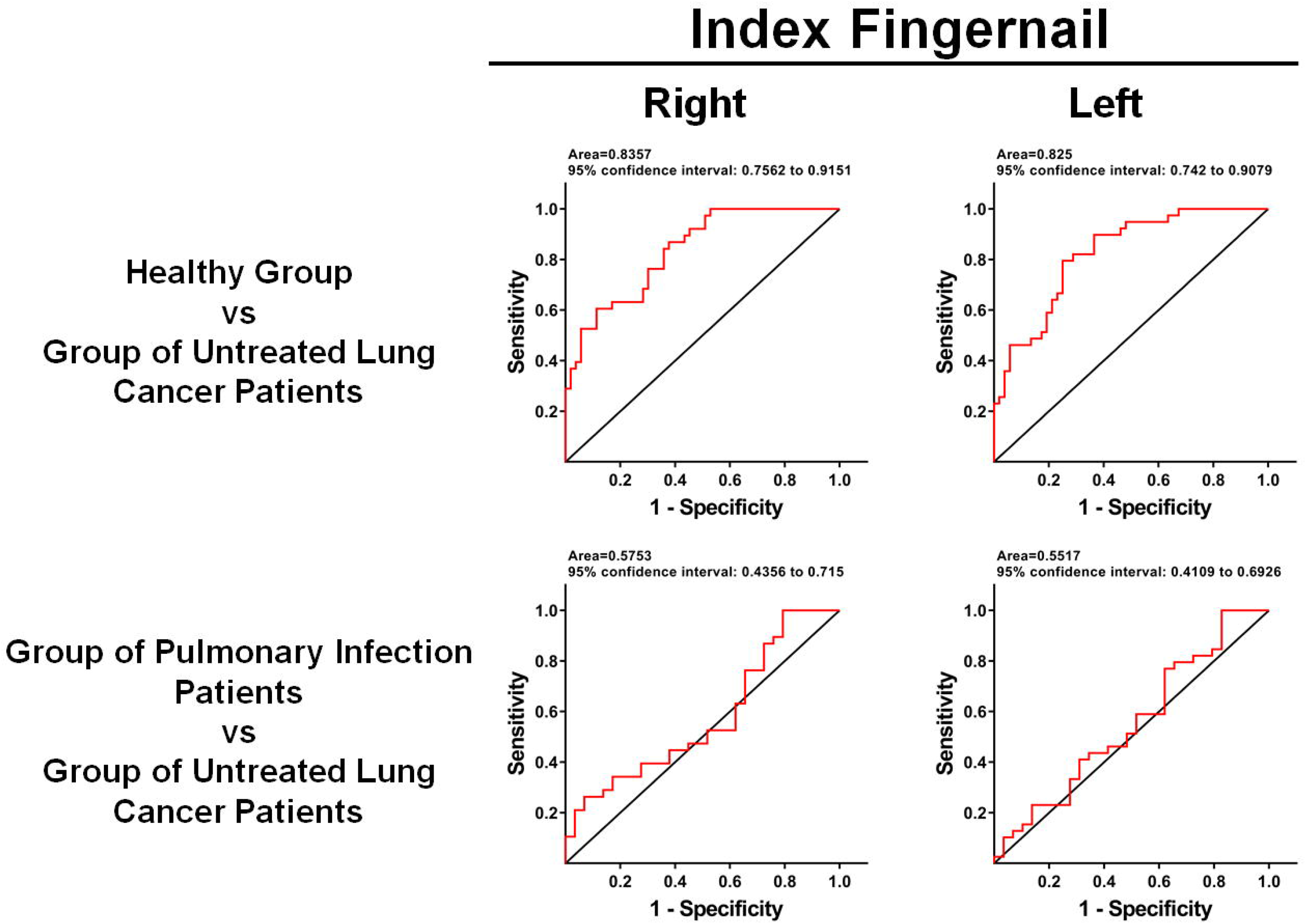
ROC analyses using the AF intensity of each examined location as the sole diagnostic parameter showed significant promise of the green AF-based technology in differentiating untreated lung cancer patients from either healthy persons or pulmonary infection patients. By using the AF intensity of each examined location as the sole diagnostic parameter, we conducted ROC analyses to determine the AUC for differentiating the healthy persons and untreated lung cancer patients (A-G). The AUC ranged from 0.8196 to 0.878 when the AF intensity of anyone of the examined locations, except right and left Ventroforefingers and left Centremetacarpus, was used as the sole diagnostic parameter. ROC analyses also showed that the AUC for differentiating the pulmonary infection patients and untreated lung cancer patients was 0.8046 and 0.8841, respectively, when the AF intensity of right or left Dorsal Centremetacarpus was used as the sole diagnostic parameter. The number of subjects in the Healthy Group, the Group of Pulmonary Infection Patients, and the Group of Untreated Lung Cancer Patients was 54, 29, and 39, respectively.

When the number of the locations with increased AF was used as the sole diagnostic parameter, our ROC analysis showed that the AUC was 0.9067 for differentiating the healthy controls and the untreated lung cancer patients (Fig. 7A). Moreover, our ROC analysis showed that the AUC was 0.7538 for differentiating the pulmonary infection patients and the untreated lung cancer patients (Fig. 7B).

**Fig. 7.**
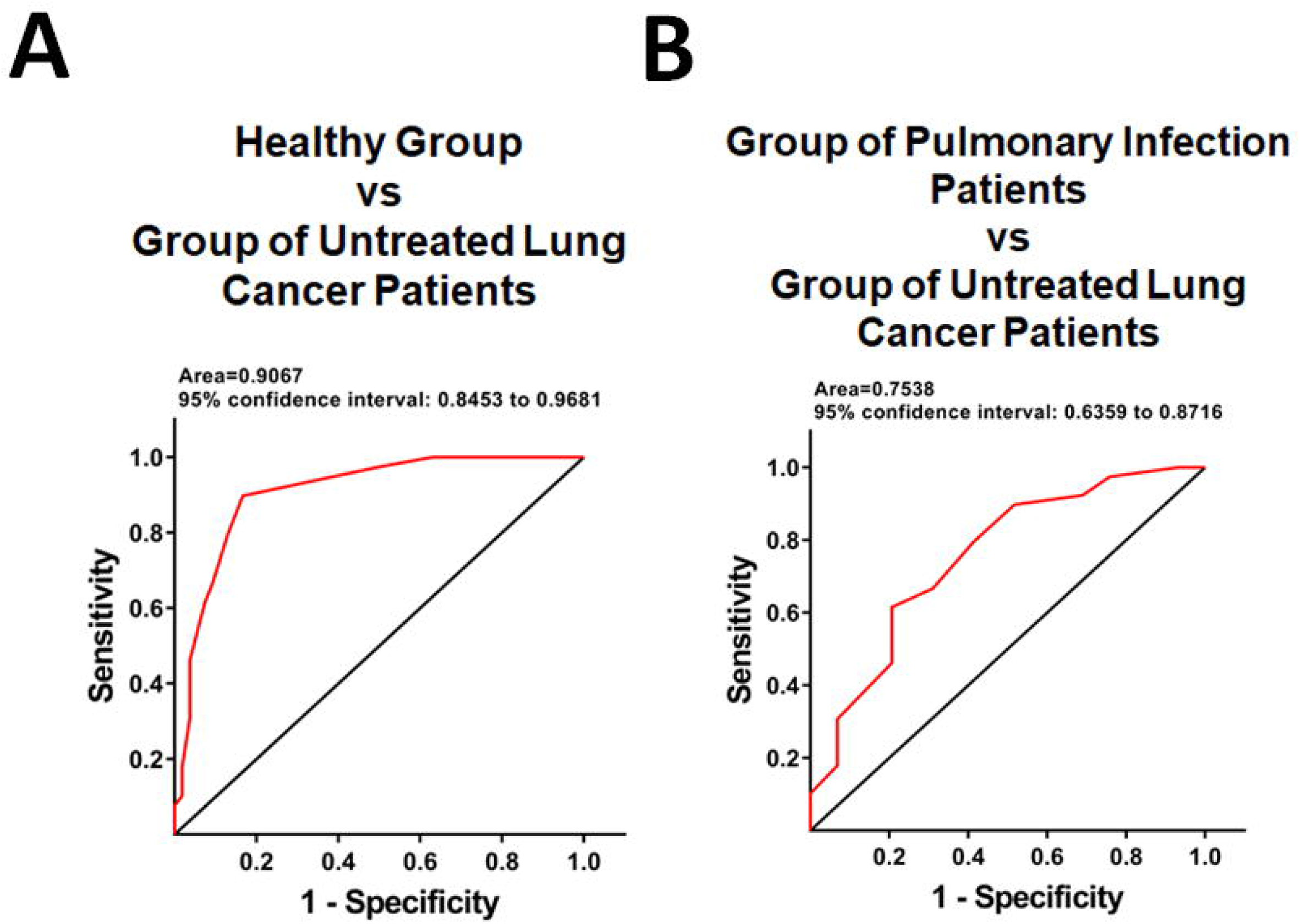
ROC analyses using the number of the locations with increased AF as the sole diagnostic parameter showed promise of the green AF-based technology in differentiating untreated lung cancer patients from either healthy persons or pulmonary infection patients. (A) When the number of the locations with increased AF was used as the sole diagnostic parameter, ROC analysis showed that the AUC was 0.9067 for differentiating the healthy controls and the untreated lung cancer patients. (B) When the number of the locations with increased AF was used as the sole diagnostic parameter, ROC analysis showed that the AUC was 0.7538 for differentiating the pulmonary infection patients and the untreated lung cancer patients. The number of subjects in the Healthy Group, the Group of Pulmonary Infection Patients, and the Group of Untreated Lung Cancer Patients was 54, 29, and 39, respectively.

### 5. The AF intensity of certain examined locations of the chemotherapy-treated lung cancer patients was significantly lower than that of the untreated lung cancer patients

We determined the green AF intensity of two index fingernails and twelve locations of the skin of the Group of Untreated Lung Cancer Patients and the Group of Chemotherapy-Treated Lung Cancer Patients. We found that the AF intensity of the right Dorsal Antebrachium, but not at other examined locations, of the chemotherapy- treated lung cancer patients was significantly lower than that of the untreated lung cancer patients (Fig. 8).

**Fig. 8.**
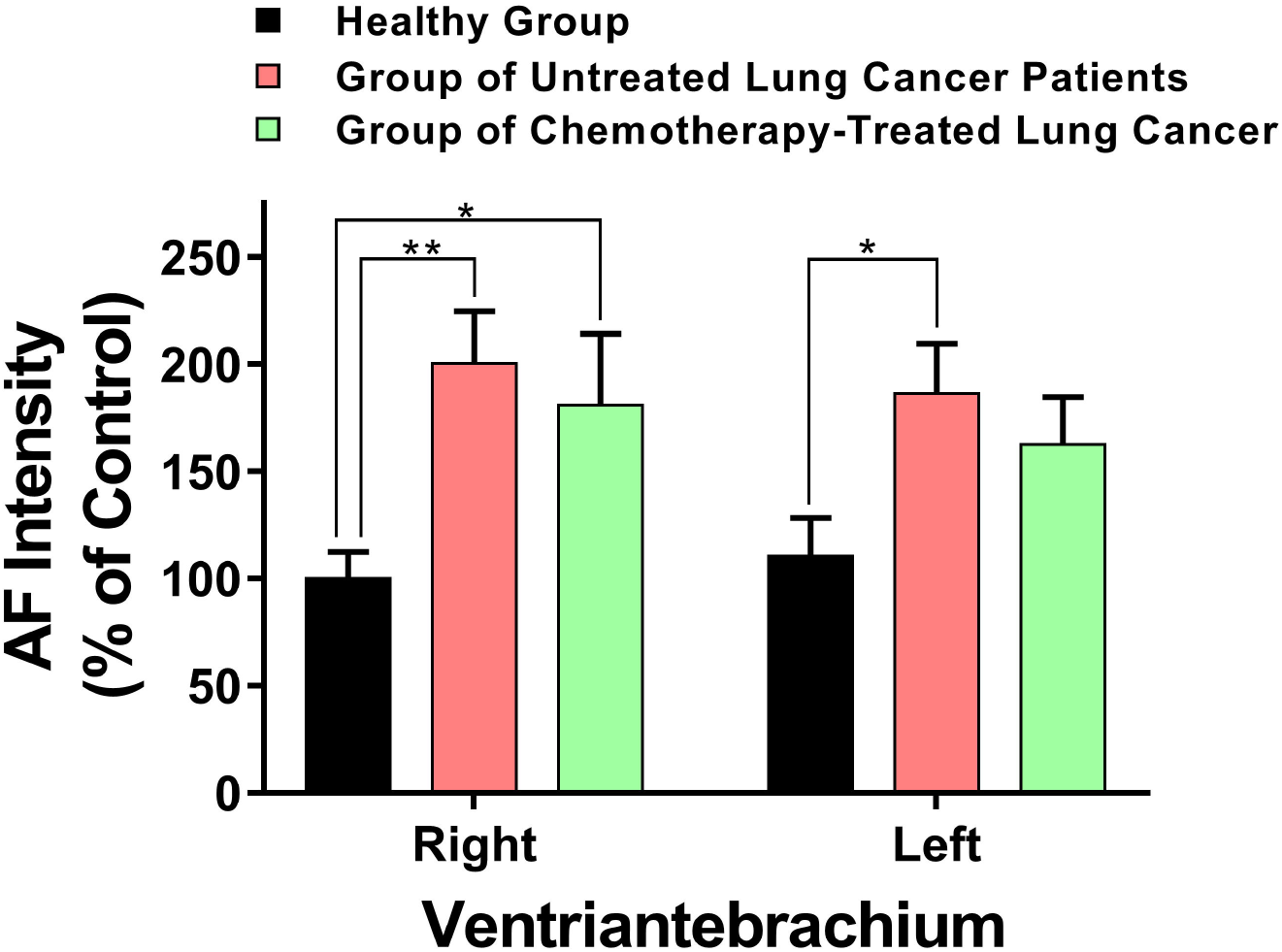
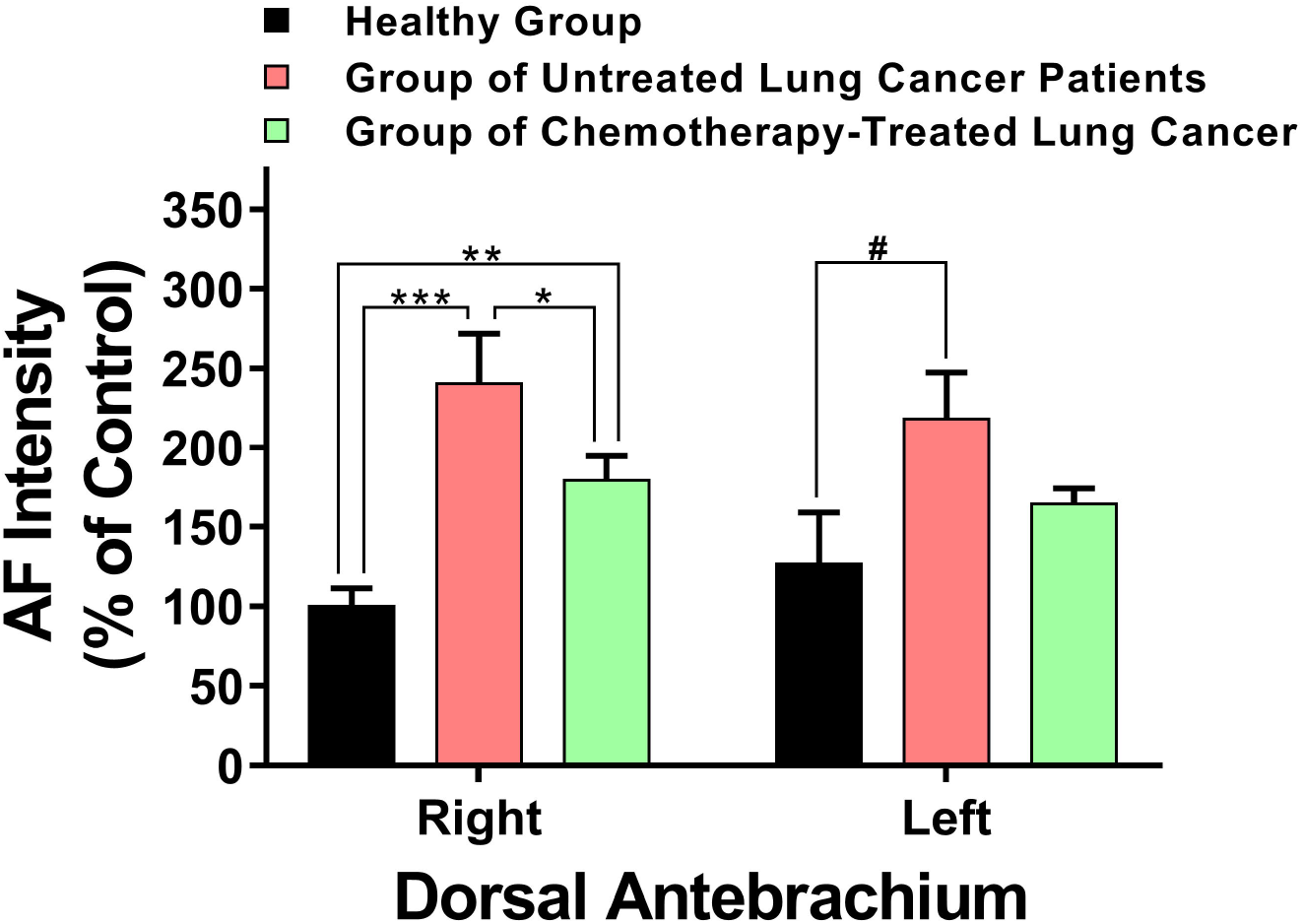
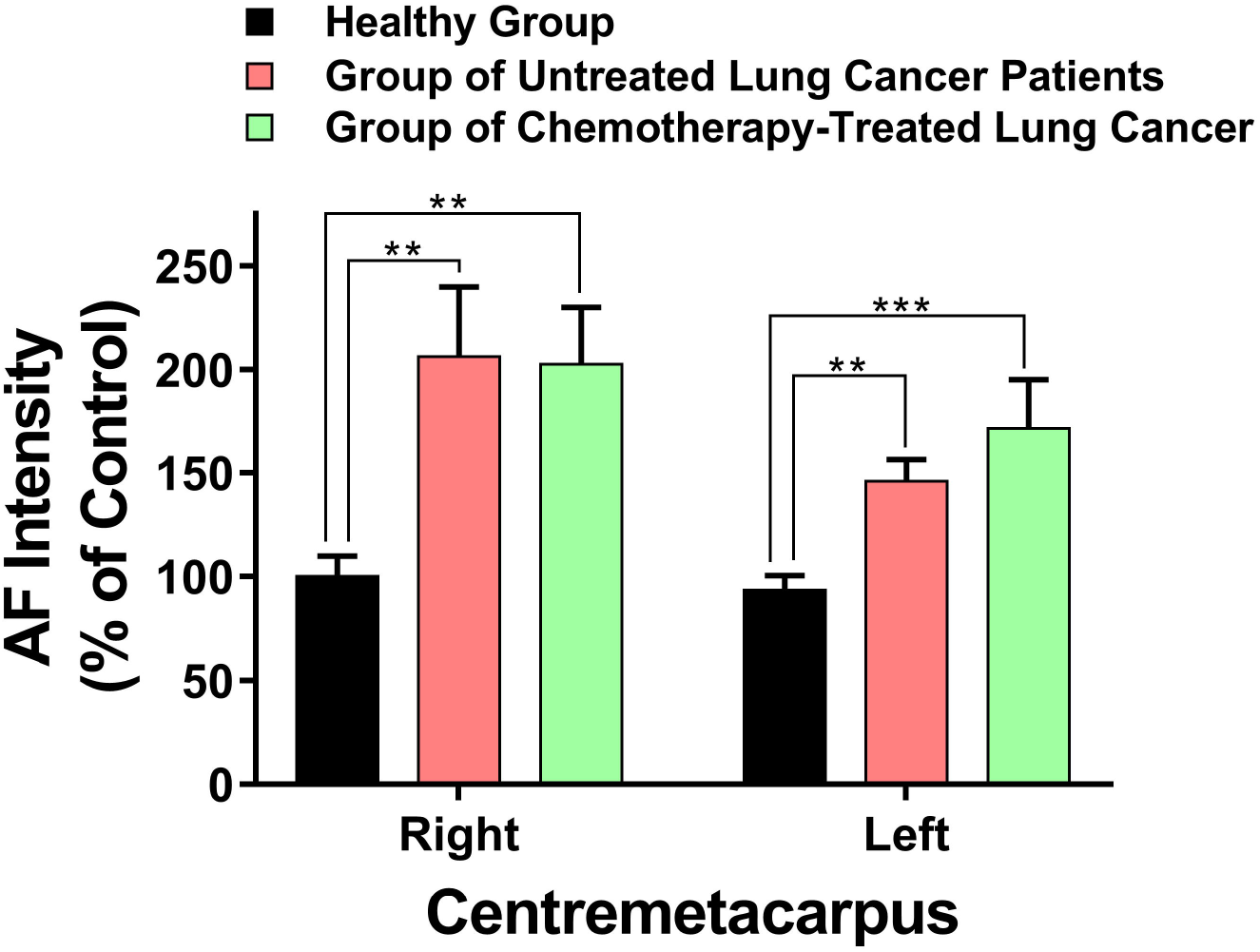
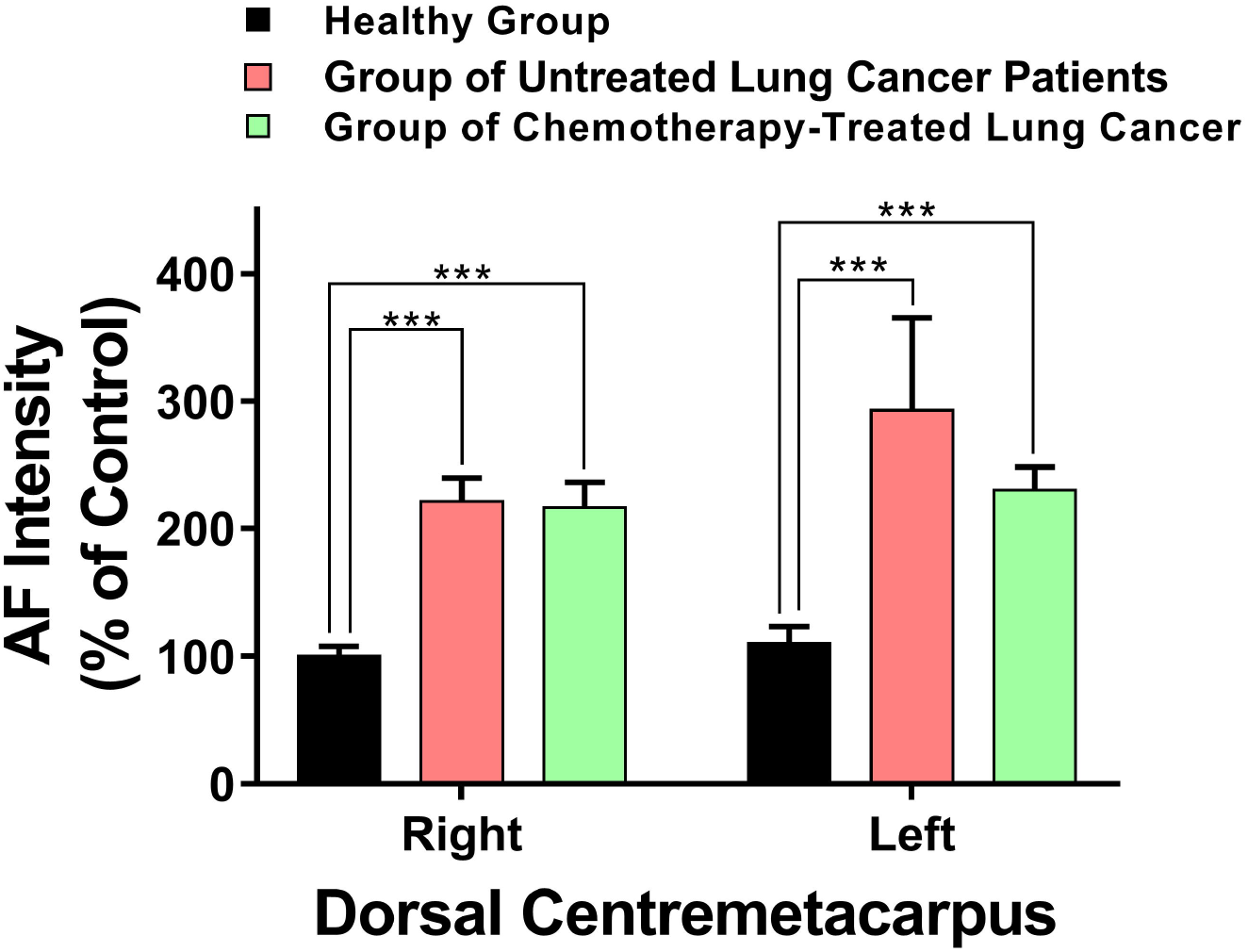
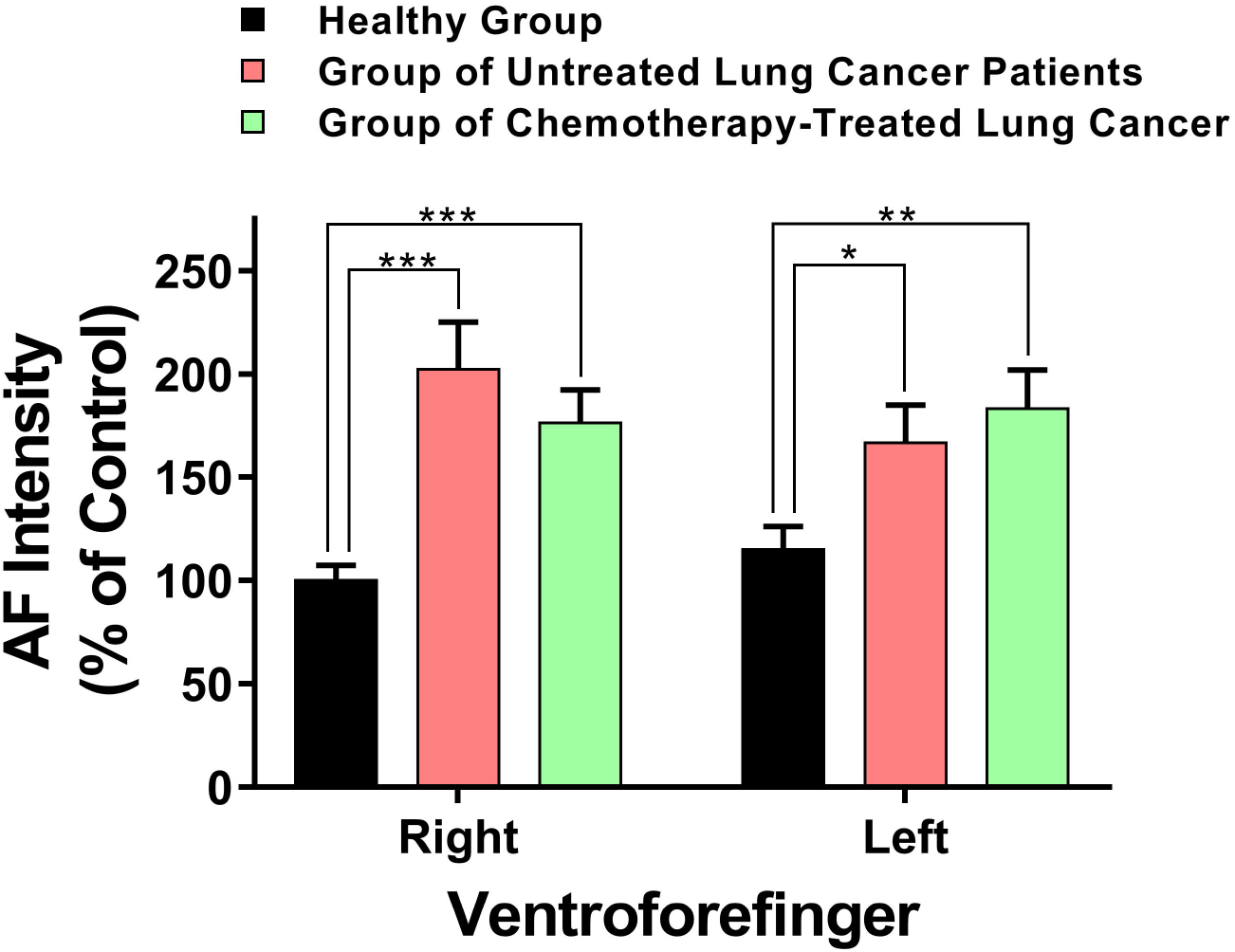
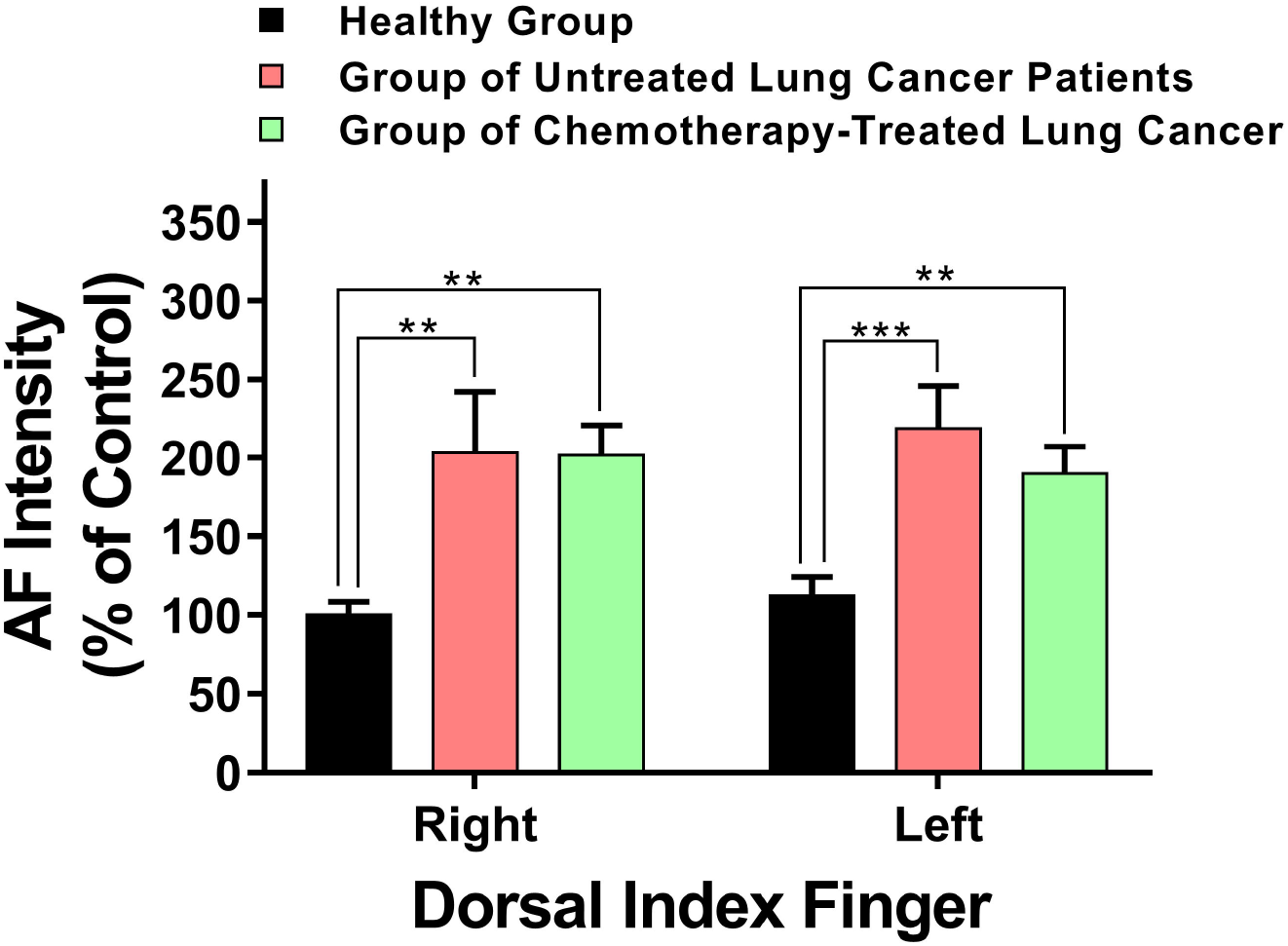
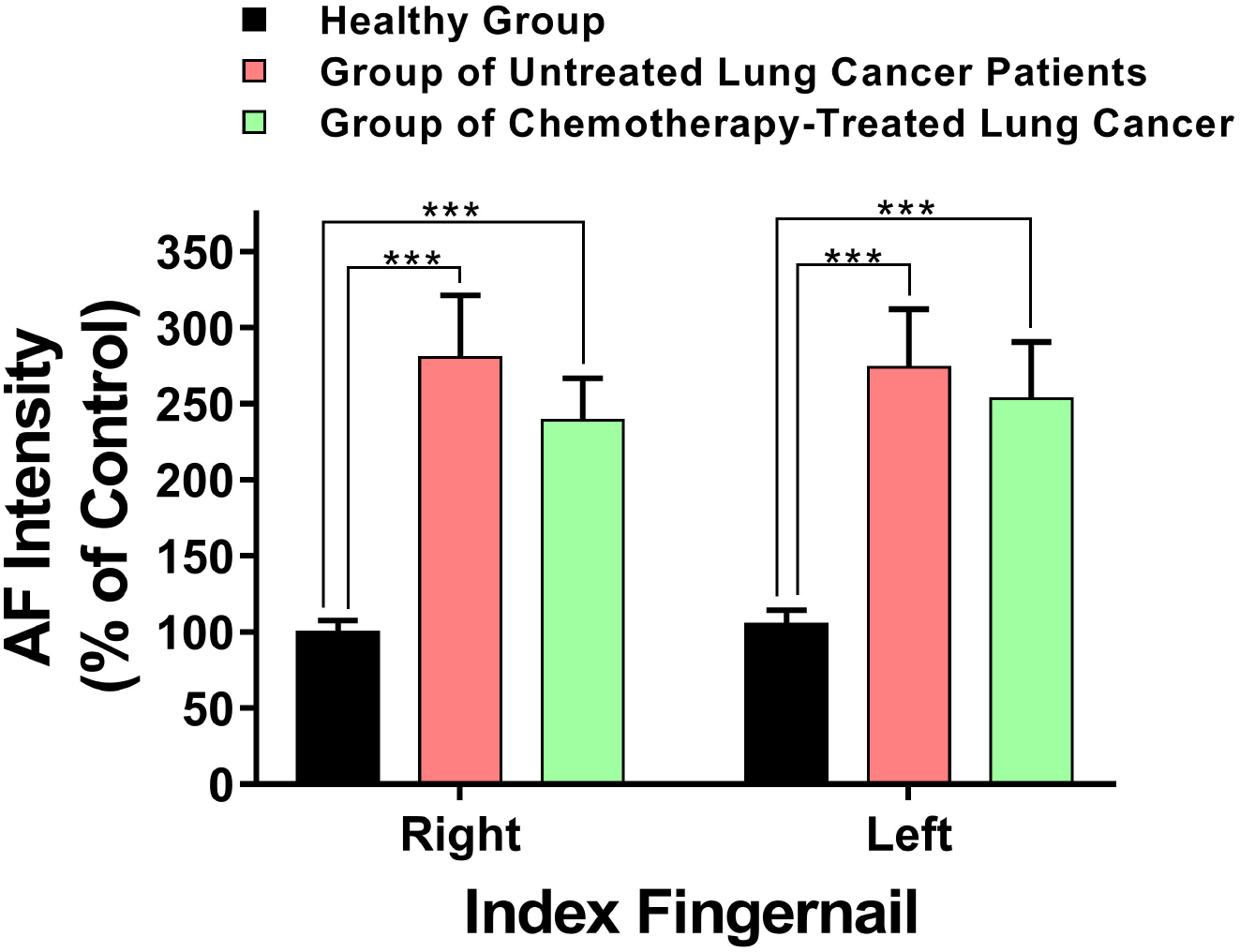
The AF intensity of certain examined locations of the chemotherapy-treated lung cancer patients was significantly lower than that of the untreated lung cancer patients. (A-G) The AF intensity of the right Dorsal Antebrachium, but not at other examined locations, of the chemotherapy-treated lung cancer patients was significantly lower than that of the untreated lung cancer patients. The number of subjects in the Healthy Group, the Group of Untreated Lung Cancer Patients and the Group of Chemotherapy-Treated Lung Cancer Patients was 54, 39, and 35, respectively.

## Discussion

The major findings of our study include: First, development of lung cancer led to a marked increase in the epidermal green AF of the mice. Second, the AF intensity of the untreated lung cancer patients was significantly higher than that of the pulmonary infection patients at certain examined locations of the skin and fingernails. Third, the ‘Pattern of AF’ of healthy controls, pulmonary infection patients and untreated lung cancer patients was markedly different from each other, e.g., in a majority of the locations examined, a higher percentage of the untreated lung cancer patients had increased AF intensity compared with pulmonary infection patients. Fourth, when the number of the locations with increased AF was used as the sole diagnostic parameter, our ROC analysis showed that the AUC was 0.9067 for differentiating the healthy controls and the untreated lung cancer patients. Fifth, the AF intensity of the right Dorsal Antebrachium of the chemotherapy-treated lung cancer patients was significantly lower than that of the untreated lung cancer patients. Collectively, our study has indicated that lung cancer patients have selective AF increases in certain locations of their skin and nails, which may become a novel diagnostic biomarker for the disease.

Early diagnosis is critical for improving the 5-year survival rate of lung cancer patients. Our current study has provided several lines of evidence suggesting that our AF-based diagnostic approach holds great promise to become a novel non-invasive diagnostic approach for lung cancer: First, development of lung cancer led to a significant increase in the epidermal green AF of the mice, indicating that it is sufficient for lung cancer to induce increased epidermal green AF of mice. Second, the AF intensity at several locations of the skin and fingernails of the untreated lung cancer patients was significantly different from that of the healthy controls and the pulmonary infection patients. Third, the ‘Pattern of AF’ of the untreated lung cancer patients was markedly different from both the ‘Pattern of AF’ of pulmonary infection patients and the ‘Pattern of AF’ of AIS patients. Fourth, when the number of the locations with increased AF was used as the sole diagnostic parameter, our ROC analysis showed that the AUC was 0.9067 for differentiating the healthy controls and the untreated lung cancer patients. Fifth, ROC analyses were conducted to determine the AUC for differentiating the pulmonary infection patients and untreated lung cancer patients, showing that the AUC was 0.8046 and 0.8841, respectively, when the AF intensity of right or left Dorsal Centremetacarpus was used as the sole diagnostic parameter.

Based on these pieces of evidence, we proposed a novel criteria for lung cancer diagnosis - the characteristic ‘Pattern of AF’ of lung cancer may be used for non- invasive, rapid and economic diagnosis of lung cancer. It is expected that the sensitivity and specificity of the ‘Pattern of AF’-based lung cancer diagnosis would be further enhanced by our following additional strategies, including addition of clinical symptoms of the patients into this diagnostic approach, applications of AI technology to conduct analyses of the structural information of the AF images, and increases of the database of the diagnostic approach. Moreover, our study has also indicated that the ‘Pattern of AF’ is a novel biomarker of lung cancer.

It is important to elucidate the mechanisms underlying the increases in the green AF of lung cancer patients. We have found that the oxidative stress induced by UVC mediates the increase in the epidermal green AF of mouse by inducing keratin 1 proteolysis (7). Our study has also shown that LPS-produced inflammation can also dose-dependently increase the epidermal green AF in mice (21). Moreover, Epidermal AF intensity was significantly associated with the serum levels of multiple cytokines in LPS-exposed mice (22). A number of studies have shown significant increases in oxidative stress and inflammation in lung cancer patients (4,6,8,14,19). Therefore, we propose that the oxidative stress and inflammation of lung cancer patients may induce increases in the epidermal AF by such pathways as inducing keratin 1 proteolysis.

Our study has also found that the AF intensity of the right Dorsal Antebrachium of the chemotherapy-treated lung cancer patients was significantly lower than that of the untreated lung cancer patients. This finding has suggested that the ‘Pattern of AF’ may also be used for evaluating the therapeutic efficacy of lung cancer.

## Acknowledgment

The authors would like to acknowledge the financial support by two research grants from a Major Special Program Grant of Shanghai Municipality (Grant # 2017SHZDZX01) (to W.Y.). and a Major Research Grants from the Scientific Committee of Shanghai Municipality #16JC1400502 (to W.Y).

## Notes

### Competing Interest Statement

The authors have declared no competing interest.

### Summary of Updates

We have added new Fig. 1 and new Fig. 2 in the revised article, which have shown our highly valuable novel findings: We used mouse model of lung cancer showing that development of lung cancer is sufficient to induce increased epidermal green AF of the mice. This finding has provided a critical base for understanding the mechanisms underlying the increased epidermal green AF of lung cancer patients.

